# Bayesian Additive Regression Trees for Genotype by Environment Interaction Models

**DOI:** 10.1101/2021.05.07.442731

**Authors:** Danilo A. Sarti, Estevão B. Prado, Alan N. Inglis, Antônia A. L. dos Santos, Catherine B. Hurley, Rafael A. Moral, Andrew C. Parnell

## Abstract

We propose a new class of models for the estimation of genotype by environment (GxE) interactions in plant-based genetics. Our approach, named AMBARTI, uses semi-parametric Bayesian additive regression trees to accurately capture marginal genotypic and environment effects along with their interaction in a cut Bayesian framework. We demonstrate that our approach is competitive or superior to similar models widely used in the literature via both simulation and a real world dataset. Furthermore, we introduce new types of visualisation to properly assess both the marginal and interactive predictions from the model. An R package that implements our approach is available at https://github.com/ebprado/ambarti.

## 1. Introduction

The interaction between genotypes and environments (GxE) is a key parameter in plant breeding (Allard and Bradshaw, 1992). Poor understanding of GxE can lead to sub-optimal selection of new genotypes and inbred lines. Understanding the GxE interactions is crucial for germplasm management, having strong genetic and economic impacts on seed production and crop yield (Sarti, 2013). Many models have been proposed for studying GxE in the context of multi-environmental experiments (METs). One special case is the additive main effects multiplicative interactions model (AMMI; Mandel, 1971).

The classical AMMI models combine features of analysis of variance (ANOVA) with a bilinear term to represent GxE interactions. Such interactions will be named here bilinear interactions. In addition, AMMI models allow for estimation of main effects of genotypes and environments as well as the decomposition of the interaction through a bilinear term. Many extensions to the AMMI models have been proposed, including robust AMMI (Rodrigues, Monteiro and Lourenço, 2016) and weighted AMMI (Sarti, 2019).

Bayesian additive regression trees (BART; Chipman et al., 2010) is a non-parametric Bayesian model that generates a set of trees and uses random splitting rules to produce predictions for a univariate response. Given its flexibility to deal with non-linear structures and richer, non-multiplicative interactions terms, the use of BART and its extensions has increased with applications in many areas including proteomic studies (Hernández et al., 2018), hospital performance evaluation (Liu et al., 2015), credit scores (Zhang and Härdle, 2010) among many others.

In this paper, we extend the AMMI model to allow for richer GxE interactions, and similarly sidestep the model choice complexity term present in all AMMI-type approaches. We achieve this goal by including a new variant of BART, which we term ‘double-grow’ BART. The new proposed method, named AMBARTI, provides a cut Bayesian model (Bayarri, Berger and Liu, 2009; Plummer, 2015), where the ‘double-grow’ BART component is solely responsible for GxE interactions.

We compare our newly proposed model with the traditional AMMI approaches and other competing interaction detection models, and we show that its performance is superior (judged on out-of-sample error) in both simulated and real-world example data. The real dataset we use is taken from the value of cultivation and usage (VCU) experiments of the Irish Department of Agriculture, which were conducted in the years between 2010 and 2019. Furthermore, the output of AMBARTI leads us to suggest several new forms of visualisation that are easier to interpret for non-specialists.

The remainder of this paper is structured as follows. In Section 2, we describe the framework used to collect evidence from METs, including the classic genetic equation to describe the relationship between phenotypes, genotypes, and environments. We also outline the formulation of the AMMI model in its classic form. In Section 2.3, we briefly describe the standard Bayesian additive regression trees model. In Section 2.4, we present the structure of our novel AMBARTI approach. Sections 3 and 4 contain the main results from the simulation experiments and real datasets, respectively. In Section 5, we conclude with a discussion and outline further opportunities. An R package that implements our approach is available at https://github.com/ebprado/ambarti (Prado and Inglis, 2022).

## 2. Methods

### 2.1. GxE interactions and MET

The phenotypic expression of a genetic character can be theoretically decomposed in terms of genetic factors, environmental factors, and the interactions between them as

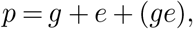

where *p* is the phenotypic response, *g* is the genetic factor, *e* is the environmental factor, and (*ge*) is the interaction between genotypes and environments. The last term is necessary due to the different response of genotypes across different environments. For instance, if we produce a rank that orders the performance of a set of genotypes into a set of environments, we will usually notice that the order of the best to worst genotype is different in each environment. The presence of GxE interactions is known to be capable of having large effects on the phenotypic response (Falconer and Mackay, 1996; Dias, 2005).

The (*ge*) terms can be estimated in a MET design where several environments and genotypes are evaluated for a given phenotype (Isik, Holland and Maltecca, 2017). In plant breeding, the need for METs is constant given the fact that the germplasm generates new genotypes every year and the pressure of diseases and other factors are dynamic. Such experiments require a complex set of logistical activities, leading to high costs of implementation. These trials thus have strong regulatory appeal in the seed and biotech industries around the world (Sarti, 2013).

Reliable information about GxE can help breeders make decisions on cultivar recommendations. Thus, models for the study of GxE need to be able to answer questions such as which genotypes can perform well across a set of environments and which are specifically recommended for a given environment. The answers to these questions are crucial both to broad breeding strategies, i.e., to obtain one or more genotypes that perform well in a set of environments, and to target breeding, where we determine the best genotype for a given environment (Sarti, 2019).

### 2.2. Traditional AMMI models

A simple statistical linear model can be used to estimate GxE effects from METs. The model can be written as

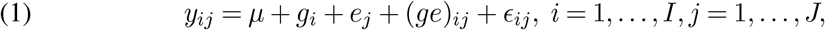

where *ε_ij_* ~ *N*(0, *σ*^2^), *y_ij_* is the phenotypic response, which represents, for example, the production of a crop in tonnes per hectare, *μ* is the grand mean, *g_i_* is the effect of genotype *i, e_j_* is the effect of environment *j*, and (*ge*)_*ij*_ represents the interaction between genotype *i* and environment *j*.

In the specification of the Equation (1), the term (*ge*) can be thought of as representing a decomposition of the residual from a more basic linear model. Gollob (1968) and Mandel (1971) proposed a method to decompose the residual term as a sum of multiplicative factors that includes the (*ge*) term. This yields the decomposition:

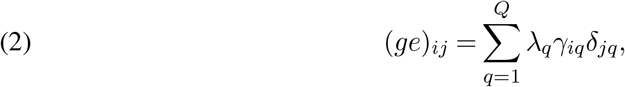

where *Q* is the number of components to be considered in the analysis, *λ_q_* is the strength of the interaction of component *q, γ_iq_* represents the importance of genotype i in component *q,* and *δ_jq_* represents the importance of environment *j* in component *q*; see the supplementary material (Sarti et al., 2022) for the restrictions imposed on *γ_iq_, δ_jq_* and *λ_q_* to make the model identifiable. Hence, the complete AMMI model is written as

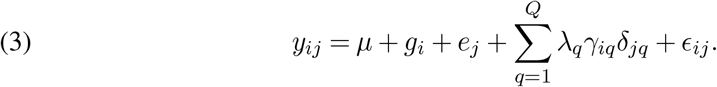

The interaction terms in (3) are estimated by a singular value decomposition (SVD) of a matrix **M**, which contains the residual values of a two-factor ANOVA model that considers the genotypes and environments as main effects. Here, *λ_q_* is the *q*-th eigenvalue of the matrix **M**, *γ_ik_* is the *i*-th element of the left singular vector and *δ_jk_* is *j*-th element of the right singular vector obtained in the SVD (Good, 1969). In practice, the classical AMMI model can be run in `R` (R Core Team, 2020) using the package `agricolae` (de Mendiburu, 2019) or via functions programmed by the user as in Onofri and Ciriciofolo (2007).

The protocol for estimation of the terms in a standard AMMI model is given by Gauch Jr (2013). This involves the following steps:

1. Obtain the grand mean and principal effects of the genotypes and environments using ANOVA with two factors based on a matrix of means containing the means of each genotype within each environment;
2. Obtain the residuals from the model above that will comprise the interaction matrix, where each row is an environment and each column a genotype;
3. Choose an appropriate value for the number of components *Q*;
4. Form the multiplicative terms that represent the reduced-dimension interactions via an SVD of the matrix of interaction residuals.

The rank of the matrix **M** is assumed to be *r* = min(*I* – 1,*J* – 1). Thus, the number of components *Q* may vary from 1,…, *r*. The quantity *r* establishes the minimum number of non-zero eigenvectors to be obtained in the SVD. Taking *Q* = *r*, the AMMI model would capture all the variance related to the interaction, and it would result in over-fitting. This problem is ameliorated by using a limited number of components *Q*. The choice of *Q* is related to the amount of total variability captured by the principal components (PCs) and, in general, is recommended to use a number of PCs that captures at least 80% of the total variability (Lal et al., 2020; Shafii and Price, 1998; Love et al., 2004; Tyagi et al., 2016; Dias and Krzanowski, 2006). Usually, the value of *Q* varies from 1 to 3.

AMMI models have been extensively used for evaluation of phenotype performance of cultivars. Nachit et al. (1992) used AMMI models to assess the performance of wheat germplasm from the International Maize and Wheat Improvement Center. Farshadfar and Sutka (2003) explored quantitative trait loci (QTL) related to adaptation in wheat. Rad et al. (2013) studied GxE for wheat in the context of drought and normal conditions. Brancourt-Hulmel and Lecomte (2003) evaluated the impact of environmental conditions in the stability of winter wheat. Badu-Apraku et al. (2012) used AMMI to study the stability of early maize genotypes in Africa. Mitroviaã et al. (2012) evaluated the performance of experimental maize hybrids using AMMI models, and Sarti (2019) studied the performance of the AMMI model in the context of simulated data. The applications of AMMI models can also be found in several other species including: a) rice (Mahalingam et al., 2006), b) barley (Mahalingam et al., 2006; Romagosa et al., 1996; Sato and Takeda, 1993; Anbessa et al., 2009), and c) sugarcane (Silveira et al., 2013).

### 2.3. Tree-based methods and BART

Introduced by Chipman et al. (2010), BART is a Bayesian model that uses a sum of trees to approximate a univariate response. In BART, each tree works as a weak learner that yields a small contribution to the final prediction. Based on a design matrix **X**, the model is able to capture interactions and non-linear relationships. The BART model can be written as

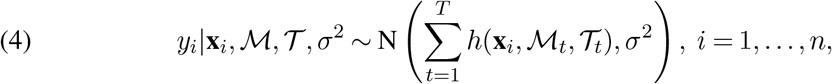

where **x**_*i*_ is the i-th row of the design matrix **X**, 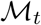 denotes the set of terminal node parameters of tree *t*, 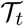 is the set of binary splitting rules that define the tree *t*, and *h*(·) = *μ_tℓ_* is a function that assigns the predicted values 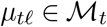 based on the design matrix **X** and tree structure 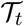. We let 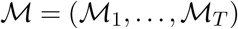 and 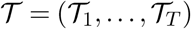 denote the sets of all predicted values and trees, respectively. Chipman et al. (2010) recommend *T* = 200 as a default for the number of trees since it works remarkably well in a wide variety of applications. However, they also suggest that *T* can be selected through cross-validation.

Unlike other tree-based methods where a loss function is optimised to grow the trees, in BART the trees are learned using a Bayesian backfitting Markov Chain Monte Carlo (MCMC) algorithm (Hastie and Tibshirani, 2000; Gamerman and Lopes, 2006; Robert and Casella, 2013). The trees are either accepted or rejected via a Metropolis-Hastings step. In addition, the trees can be learned by four moves: grow, prune, change or swap. It is important to highlight that in all moves the splitting rule is defined by randomly selecting one covariate and one split point. In the grow move, a terminal node is selected and then two children nodes are created below it. When pruning, a parent of two terminal nodes is selected and its children nodes are removed. During the change move, a parent node is picked and its splitting rule (i.e., covariate and split point) is redefined. In the swap move, a pair of parent-child internal nodes is chosen and the splitting rules associated to the two nodes are swapped.

As a fully Bayesian model, BART assumes prior distributions on all quantities of interest. First, the node-level parameters *μ_tℓ_* are assumed to be i.i.d 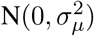, where 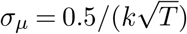 and *k* ∈ [1,3]. Second, the residual variance *σ*^2^ is assumed to be distributed as *IG*(*ν*/2, *νλ*/2), where IG(·) denotes an Inverse Gamma distribution. Third, to control how shallow/deep a tree may be, each non-terminal node has a prior probability of *α*(1 + *d*)^*β*^ of being observed, where *α* ∈ (0,1), *β* ≥ 0, and d corresponds to the depth of the node; Chipman et al. (2010) recommends *α* = 0.95 and *β* = 2 as default values. These hyperparameter values tend to select trees which are not too deep.

We remark that the priors above are a crucial element to identify *σ*^2^ as they control the tree topology and the variability of the prediction at the terminal node level. Were these priors to be set too vague (e.g., so that deep trees were favoured), then the value of *σ*^2^ would shrink towards zero and we would be left with an over-fitted model. Thus, these priors force the trees to be shallow and shrink the terminal node parameters towards zero such that each tree only explains a small component of the data, which leaves some variation in the residuals and resolves the identifiability issue of *σ*^2^.

The identification of the interaction effects in BART is powered by the full conditional of the trees, which we denote by 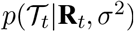, where 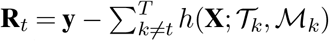 represents the vector of the partial residuals excluding tree *t;* see the supplementary material (Sarti et al., 2022) for details. As in BART the trees are learned by using splitting rules that are created by randomly selecting a covariate and a split point, 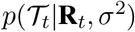 is utilised to select only ‘good’ trees (i.e., trees that help reduce the residual variance).

Finally, the structure of the BART model for a continuous response can be summarised as follows. First, all 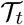 are initialised as stumps. Then, each tree is learned, one at a time, using one of the four moves previously described (grow, prune, change or swap). Next, the newly proposed tree 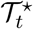 is compared to its previous version 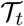 via a Metropolis-Hastings step taking into account the partial residuals **R**_*t*_ and the structure/depth of 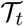 and 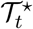. After that, the predicted values for each terminal node *ℓ* of the tree *t* are generated and then *σ*^2^ is updated. For a binary outcome, the data augmentation strategy of Albert and Chib (1993) can be used; see Tan and Roy (2019) and Prado et al. (2021) for more details.

Due to its flexibility and excellent performance on regression and classification problems, BART has been applied and extended to credit modelling (Zhang and Härdle, 2010), survival analysis (Sparapani et al., 2016; Linero et al., 2021; Basak et al., 2021), proteomic biomarker analysis (Hernández et al., 2015), polychotomous response (Kindo et al., 2016) and large datasets (Hernández et al., 2018; Linero and Yang, 2018). More recently, papers exploring the theoretical aspects of BART have been developed by Jeong and Ročková (2020); Ročková and van der Pas (2020); Ročková and Saha (2019); Linero and Yang (2018).

### 2.4. AMBARTI

To insert the BART model inside an AMMI approach, we make some fundamental changes to the way the trees are learned and structured. As a first step, we can write the sum of trees inside the Bayesian version of the AMMI model as

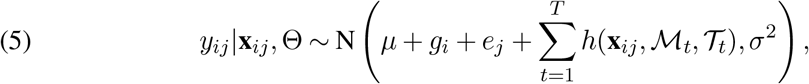

where *y_ij_* denotes the response for genotype *i* and environment 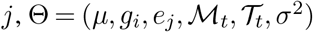, *μ* is the grand mean, and *g_i_* and *e_j_* denote the effects of genotypes and environments, respectively. The component 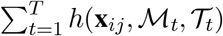 is the same as presented in (4) and **x**_*ij*_ contains dummy variables that represent combinations of *g_i_* and *e_j_*. In order to get the posterior distribution of the new parameters, we assume that 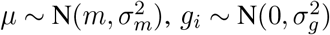 and 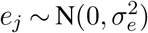 as well as that 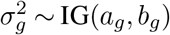 and 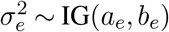.

At first look our model is similar to the semi-parametric BART proposed by Zeldow et al. (2019). However, our approach differs in that i) we do not partition the covariates into two distinct subsets, as the dummy variables (*g_i_* and *e_j_*) that are used in the linear predictor are also contained in **x**_*i*_; ii) most importantly, we add to the tree-generation process in BART a ‘double grow’ and a ‘double prune’ steps so that we guarantee that the trees will include at least one *g_i_* and one *e_j_* as splitting criteria and; iii) unlike Zeldow et al. (2019), we do not use the full residuals 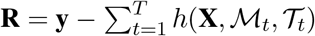 to update the linear predictor estimates, but rather the response variable only.

The rationale of the double grow and double prune moves is to force the trees to exclusively work on the interactions between *g_i_* and *e_j_*. In doing so, we remove the chance that the ‘single’ moves split on a single *g_i_* or *e_j_* variable, which would lead to confounding with the main marginal genomic or environment effects. For example, in the double grow, rather than randomly selecting one covariate and one split point when growing a stump, a variable *g*✶ is chosen and then another variable *e*✶ is randomly selected and both define the splitting rules of the corresponding tree. Here, the dummy variables *g*✶ and *e*✶ are sampled from the sets of all possible combinations of g**¿** and *e_j_*, respectively. Conversely, in the double prune move a tree is pruned twice to prevent it from having a single *g_i_* or *e_j_*. Thus, the resulting tree from this double move will always be a stump.

We allow for extra flexibility in the identification of the interaction effects by setting up the design matrix of the BART model to contain all possible combinations of genotype and environment effects. For instance, when *I* = 5 genotypes and *J* = 6 environments, 2^*I*–1^ – 1 = 15 and 2^*J*–1^ – 1 = 31 unique dummy variables are generated for *g* and *e*, respectively; see Wright and König (2019) for splitting methods on categorical covariates. In this case, there are 46 dummy variables available so that BART can (double) split on them to identify possible interaction between genotypes and environments. To illustrate this, in Figure 1 a dummy variable is sampled from the set of 15 dummy variables related to *g* along with a dummy variable from the set of 31 dummy variables associated with *e*.

**Fig 1:**
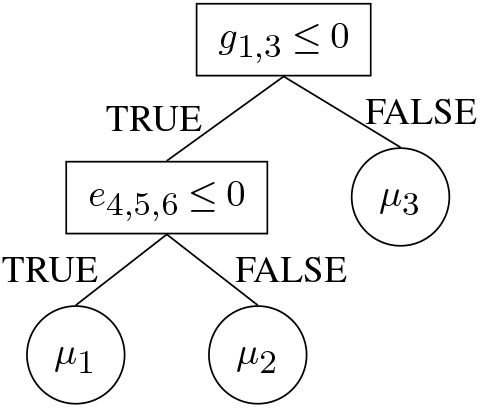
An example of a tree generated from a double grow move based on the sets of dummy variables generated for *g* and *e*. *g*_1,3_ denotes a dummy variable that combines genotypes 1 and 3, and *e*_4,5,6_ represents a dummy variable that binds environments 4, 5, and 6.

As suggested by a referee, an alternative to what we did is i) to use *k* categorical variables over the set of 2^*k*-1^ – 1 combinations and also ii) modify the prior on the trees so that it would not overly penalise deep trees. The first approach is used in the current `R` packages that offer implementations of BART, such as `bartMachine` (Kapelner and Bleich, 2016), `BART` (McCulloch et al., 2020) and `dbarts` (Dorie, 2020), which all represent categorical variables with *k* > 2 levels into *k* dummy variables. In contrast, in random forests the standard approach to deal with categorical variables is to create all the 2^*k*-1^ – 1 2-partition combinations (Wright and König, 2019) subject to a not too big *k*. However, we explored the two alternatives mentioned above and the results with k dummy variables were not encouraging, whilst the change to the set of 2^*k*-1^ — 1 combinations vastly improved the prediction performance.

We recall that before estimating the parameters in the bilinear term of the AMMI model in (3), the effects of *g_i_* and *e_j_* are estimated via a linear regression. Hence, the residuals are organised into an *I* × *J* matrix, in which the rows and columns sum to zero, and then an SVD is performed on the residual matrix. Due to the sum-to-zero constraint, the resulting output from the bilinear term sums to zero within each *g_i_* and *e_j_*. In a similar fashion, a ‘fully Bayesian AMBARTI’ would have to guarantee that the predictions from the BART component sum to zero within the *g_i_* and *e_j_* in order to be comparable to AMMI. However, if we impose the sum-to-zero constraint on the BART predictions within the *g_i_* and *e_j_*, this is equivalent to cutting feedback (i.e., remove without marginalisation) the BART predictions from the full conditionals of *g_i_* and *e_j_*; see the supplementary material (Sarti et al., 2022) on the AMBARTI algorithm. In Section 4, we compare AMBARTI with a set of comparators that includes a fully Bayesian AMMI model, which performs poorly compared to AMBARTI and other interactions models.

The motivation behind cutting feedback in Bayesian models is studied in Bayarri, Berger and Liu (2009) and Plummer (2015). In the first work, the authors list a set of situations where the ‘modularisation’ (i.e., cut feedback) of Bayesian models can be useful. They motivate the use of modularisation through examples in the analysis of computer models and present possible reasons for performing it. They point out that Bayesian models that eventually incur identifiability/confounding issues might require modularisation, especially when there is interest in determining effects of interest and not only the overall prediction. We highlight that were AMBARTI fully Bayesian rather than a cut Bayesian model, there would be some bias/identifiability issues, due to the sum-to-zero condition mentioned above, between the *g_i_* and *e_j_* and the BART component. Finally, Plummer (2015) highlights some issues that can affect the sampling of the parameters from a cut Bayesian model and also proposes solutions to circumvent some of the issues. In Section 2.4.1, we tackle Plummer’s concerns by showing that the posterior samples from AMBARTI are valid.

An appealing advantage of AMBARTI over AMMI is that it does not require the specification of the number of components *Q* in the bilinear sum and does not require complex orthonormality constraints on the interaction structure; see the supplementary material (Sarti et al., 2022) for the constraints of the AMMI model. In a Bayesian context, these constraints can lead to complex prior distribution choices for implementation of AMMI (as in Josse et al., 2014; Crossa et al., 2011). Furthermore, although AMBARTI adds a computational cost to the BART model, we have found this to be negligible for standard MET datasets that usually have values of *I* and *J* up to the low tens or hundreds.

An additional advantage of using a (cut) Bayesian approach as in AMBARTI is that we have access to the posterior distribution of each parameter. As the model is fitted, we are thus able to ascertain the general levels of uncertainty in each *g_i_* or *e_j_* component, which may assist with future experimental designs. Similarly, the interaction term is also estimated probabilistically, and so may avoid interpretation errors associated with, e.g., biplots from a traditional AMMI model.

In terms of estimation, the AMBARTI model can be fitted as follows. First, the parameter estimates *g_i_* and *e_j_* are sampled taking into account the response variable **y** (not the residuals). Then, one at a time, the trees are updated via partial residuals 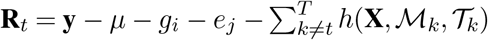. Hence, the terminal node parameters are generated and the sample variance is updated. In the end, posterior samples associated with 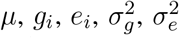, and 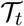 are available, which allow for the calculation of credible intervals and evaluation of the significance of the parameter estimates; see the supplementary material (Sarti et al., 2022) on the AMBARTI algorithm.

#### 2.4.1. Validity of the posterior sampling in AMBARTI

The AMBARTI model estimates the genotype and environment main effects taking into account only the response and disregarding the BART component. In contrast, when fitting the BART component, the effects of *g* and e are taken into account. We set up AMBARTI in this way as we aimed to compare its results to the frequentist AMMI model. We recall that in classical AMMI the estimation of g and e is first carried out and then the interactions between them are estimated without the bilinear term being fed back into the process to update the estimates of g and e. As stated, AMBARTI can be seen as a modularised model (Bayarri, Berger and Liu, 2009) or a cut Bayesian model which uses the ‘naive cut algorithm’ (Plummer, 2015). Nonetheless, we show that our model, under the naive cut algorithm, generates valid posterior samples following Plummer (2015).

A simple way of writing our model, given the tree structures, is:

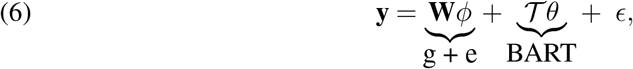

where 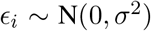, **W** is a binary matrix of values allocating **y** to the correct genotype/environment, *ϕ* are the *g* and e parameters, 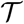 is a matrix that relates each of the trees to the individual observations, and *θ* are parameters of the interactions. Under this formulation, we have that the quantities of interest *ϕ*, 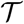 and *θ* can be estimated via their full conditional distributions:

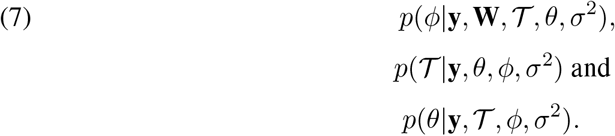

Figure 2 presents AMBARTI from the perspective of a cut Bayesian model. The likelihood function depends on *θ*, 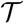 and *ϕ*, and all parameters are of interest. The graph is divided in two parts (*G*_1_ and *G*_2_). The idea of the naive cut algorithm is that when constructing the full conditionals for parameters in *G*_1_-likelihood terms involving random variables in *G*_2_ are ignored (Plummer, 2015).

**Fig 2:**
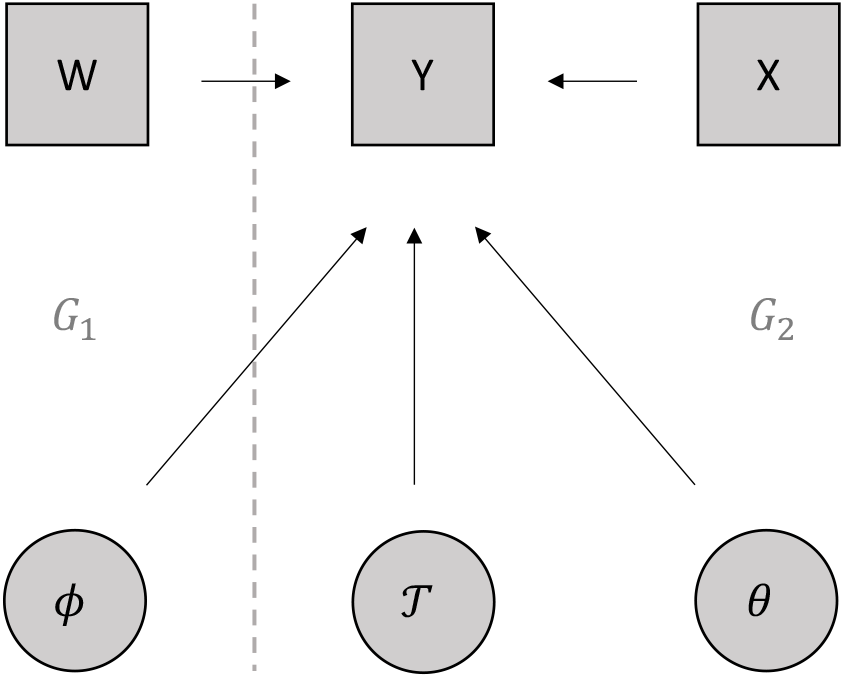
Graphical representation of the AMBARTI model.

The ‘cut’ conditional distributions for *ϕ* and *θ* are given by:

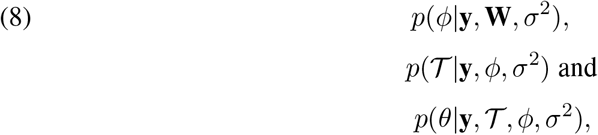

where 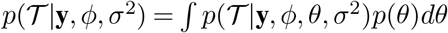. The full conditional in (7) considers the *full* Bayesian model, whilst in (8) there is no dependence/influence of 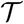 and *θ* over *ϕ*. Plummer (2015) describes three situations when the feedback from one component may not be helpful to the other. First, when the relation between the response and *θ* is speculative. Second, when there is conflict between the sources such that 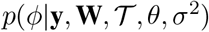 is very different from *p*(*ϕ*/**y, W**, *σ*^2^). Third, when there are computational issues in terms of convergence and mixing.

Plummer (2015) states that the cut in the feedback of 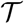 and *θ* into the conditional distribution for *ϕ* may lead to lack of convergence to the (cut) posterior distribution. However, for the cut algorithm to draw approximate samples from the cut model, one of the two conditions below need to be verified.

1. The transition from *ϕ* to *ϕ*’, 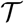 to 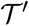 and *σ*^2^ to *σ*^2’^ cannot cause significant changes in the conditional distribution for 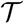 and *θ*, i.e., the limit of

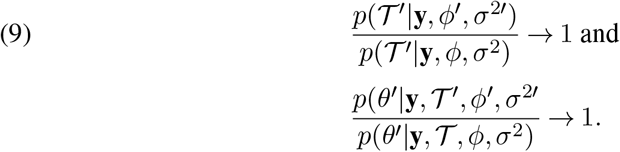 Under the cut Bayesian model framework, this condition makes sense to parameters that do not have closed-form full conditionals (e.g., the full conditional for the tree in AMBARTI). In an attempt to satisfy this condition, Plummer (2015) proposes an algorithm based on tempered transitions which only allows small steps for *ϕ*, 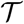 and *σ*^2^.
2. The probabilities of moving from 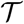 to 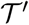 and *θ* to *θ*’, denoted by 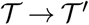 and *θ* → *θ*’, do not depend on 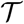 and *θ*, respectively, i.e., the limit of

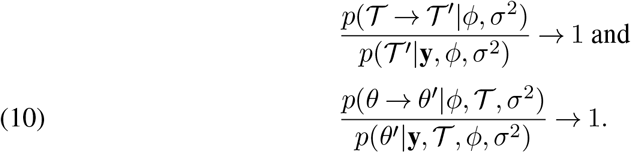 We remark that all parameters in the AMBARTI model attain the condition in (10), except 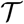. For example, the probability of *θ* → *θ*′ depends exclusively on the posterior conditional distribution *p*(*θ*’|**y, X**, *ϕ*, *σ*^2^), which in turn does not depend on the previous *θ* since it has closed-form. To make our point clearer, recall expression (6) and that in BART, *a priori*, 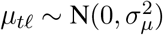, which is completely specified with no dependence on its previous values. In addition, the posterior conditional distributions of *g_i_, e_j_, μ_tℓ_*, and *σ*^2^ have all closed-form expressions, which are presented in the supplementary material (Sarti et al., 2022). In relation to the transition probability of an individual tree 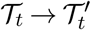, we point out that it does rely upon the previous tree 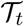 since the transition kernel 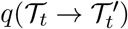, specifically for the grow and prune moves, depends on the number of terminal and internal nodes of 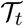; see Appendix A of Kapelner and Bleich (2016) for further details on the transition probabilities of the moves in BART. However, we empirically show, in the supplementary material (Sarti et al., 2022), that the transition 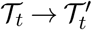 under the naive and tempered cut algorithms do not differ, which supports the condition in (9) for AMBARTI.

## 3. Simulation Study

In this Section, we compare AMMI, the Bayesian version of AMMI (B-AMMI) proposed by Josse et al. (2014), and AMBARTI using the root mean square error (RMSE) for predicted values *y* and for the interaction term on out-of-sample data. Our simulation experiment was carried out considering two scenarios. In the first, we simulated data from the AMMI model with *Q* = {1,2,3} and then fitted AMBARTI, B-AMMI, and AMMI. In the second scenario, we simulated data from the AMBARTI model and then fitted AMBARTI and three AMMI (and B-AMMI) models with different number of components to describe the interactions (i.e., *Q* = {1,2,3}). In both scenarios, we fitted the models to a training set with *I* × *J* observations and evaluated the performance on an out-of-sample test set of the same size.

For both scenarios, we set *I* = *J* = {10,25}, *μ* = 100 and generated *g_i_* and *e_j_* from 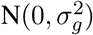 and 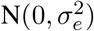, respectively, where *σ_g_* = *σ_e_* = {1,5}. The parameters *γ_ik_* and *δ_jk_* were generated from N(0,1) and then the orthornormality constraints were applied following the results presented in the supplementary material (Sarti et al., 2022). In addition, for *Q* = 1, we consider two values for *λ* (i.e., *λ* = {8,12}); for *Q* = 2, *λ* = ({12,8}, {12,10}) and; for *Q* = 3, *λ* = {12,10,8}. In the simulation from AMBARTI, we set *T* = 200 trees and generated each tree by using the ‘double grow’ move considering 2^*I*–1^ – 1 possible covariates for *g_i_* and 2^*J*–1^ – 1 for *e_j_*.

Finally, the AMMI model used in the simulations is presented in Equation (3), which represents a completely randomised trial design. The AMBARTI model used is shown in Equation (5).

### 3.1. Simulation results

We start simulating synthetic data from the AMMI system, which is the harshest test for the AMBARTI model. Figure 3 shows the RMSE values for *ŷ* based on the out-of-sample sets of three models considering 10 Monte Carlo repetitions. The datasets considered in this Figure were simulated considering *I* = 10 genotypes and *J* = 10 environments, with different values of *Q* = {1,2,3}, and two values for the genotypic and environmental standard errors *σ_g_* = {1,5} and *σ_e_* = {1,5}, respectively.

**Fig 3:**
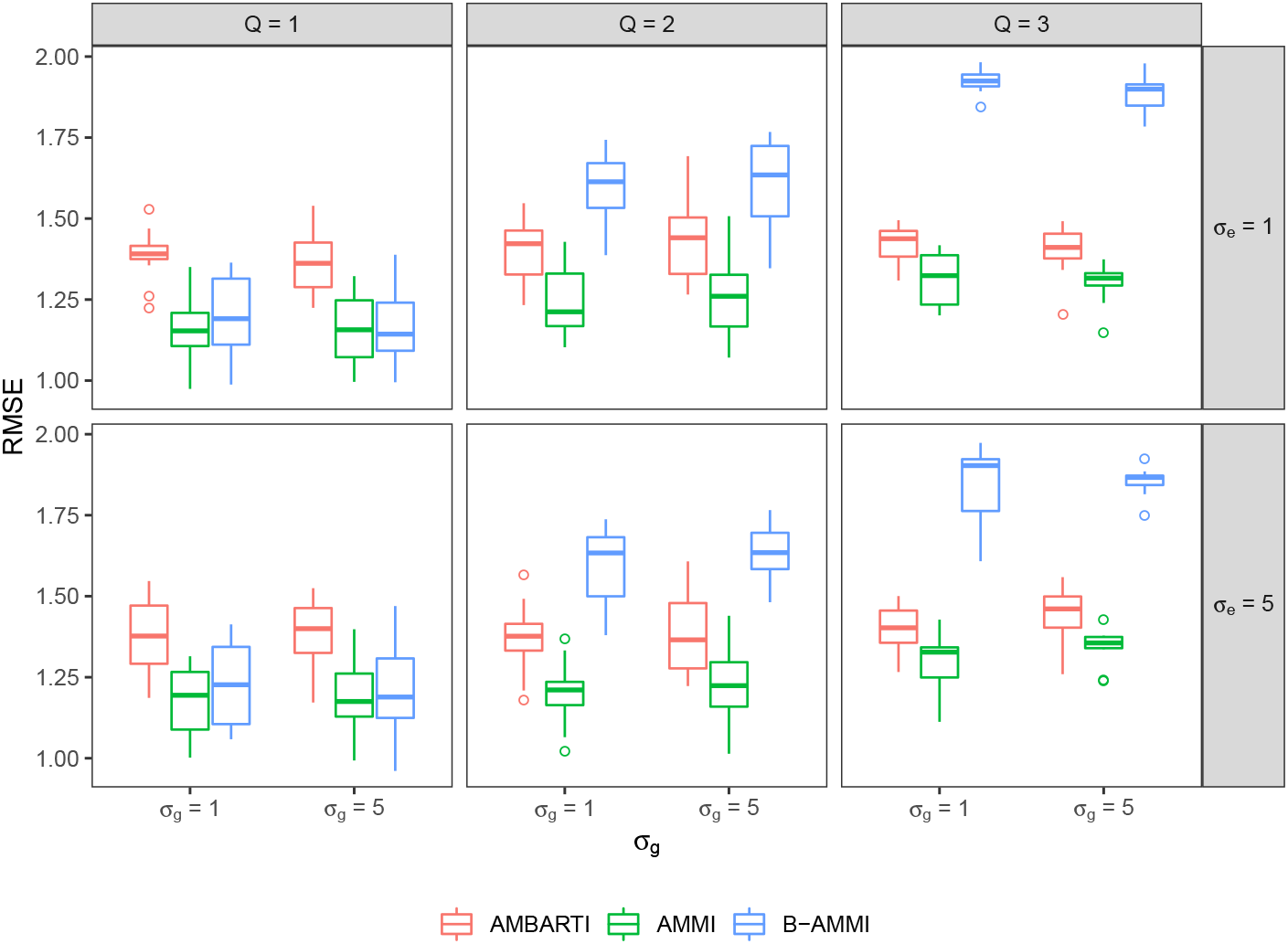
Out-of-sample RMSE for *ŷ* based on the results of AMMI, B-AMMI, and AMBARTI for data simulated from the AMMI model with *I* = *J* = 10. The different panels contain 10 Monte Carlo repetitions and represent different combinations of the simulated parameters for the creation of the dataset. Unsurprisingly, AMMI performs very well here, with AMBARTI having RMSE values around 17% higher.

As the data were simulated from the AMMI equation, we would expect that the AMMI model would perform exceedingly well, and this is what we see in general considering all the results of Figure 3. More specifically, we can see in the first upper panel that AMBARTI has higher RMSEs compared to AMMI for all values of *Q*. Further, we see that B-AMMI has similar performance to the frequentist AMMI only for *Q* = 1. As the number of components in the bilinear term increases, the results from B-AMMI deteriorate compared to AMMI and AMBARTI. In addition, it is possible to note that there is no clear effect of *σ_g_* or *σ_e_* on the RMSEs. However, even with the AMMI model presenting the best results, AMBARTI demonstrates highly competitive performance, with RMSE values around 17% higher than that of the AMMI model.

Figure 4 shows the results of the second simulation scenario, where the data were simulated from the AMBARTI equation. Again, different combinations of parameters were used in the simulation of the training and out-of-sample sets. The upper panels show results for *I* = 10 genotypes and *J* = 10 environments; the lower ones for *I* = 25 and *J* = 25. Furthermore, three AMMI and three B-AMMI models were fitted considering *Q* from 1 to 3. In this case, the AMMI model, even with high values of *Q*, performs very poorly with RMSE values 3 times higher on average than that of AMBARTI for *I* = 10. For *I* = 25, the AMMI model with *Q* = 3 is competitive with AMBARTI, with the latter being slightly better. In addition, we can see that AMBARTI presents better results when compared with B-AMMI, regardless of the value of *Q*. In this comparison, it is worth mentioning that more complex possibilities of interactions may be obtained when simulating from AMBARTI compared to AMMI.

**Fig 4:**
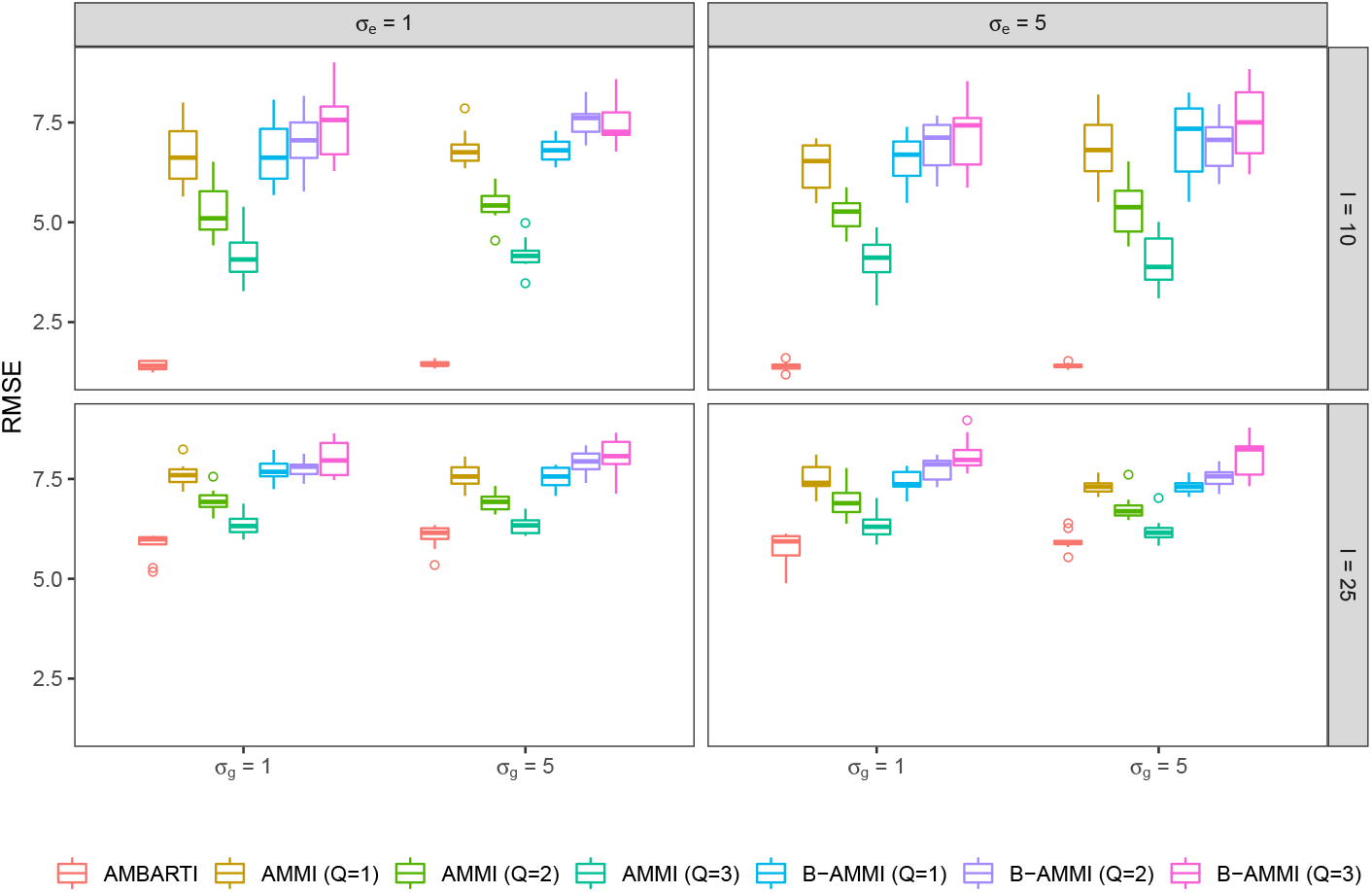
Out-of-sample RMSE for *ŷ* based on the results of AMMI (with varying *Q*), B-AMMI, and AMBARTI for data simulated from the AMBARTI model with *I* = *J* = 10 and *I* = *J* = 25. The boxplots contain 10 Monte Carlo repetitions. The AMMI RMSE values are on average 3 times higher than that of AMBARTI for *I* = 10.

The next important comparison to be made between AMBARTI and AMMI is related to the interaction term (i.e., the bilinear term for AMMI and the BART component for AMBARTI). Such tests are shown in Figures 5 and 6, where we show the RMSE performance only for the interaction component.

**Fig 5:**
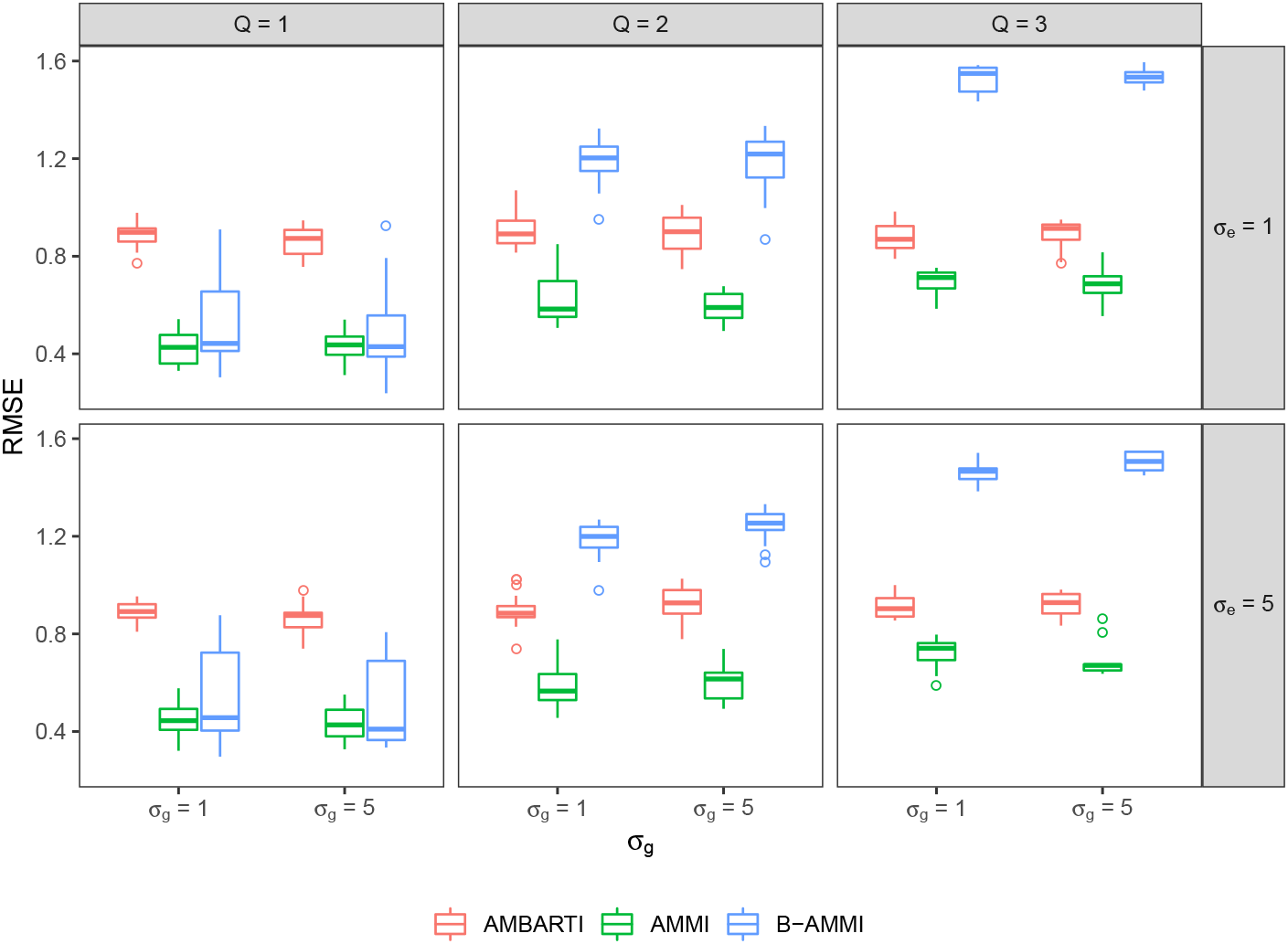
Out-of-sample RMSE related to the interaction term of AMMI models for data simulated from AMMI. The different panels show the different parameter values used in the simulation. The performance of AMMI here is optimal, with AMBARTI performing slightly worse than AMMI when *Q* = 3.

**Fig 6:**
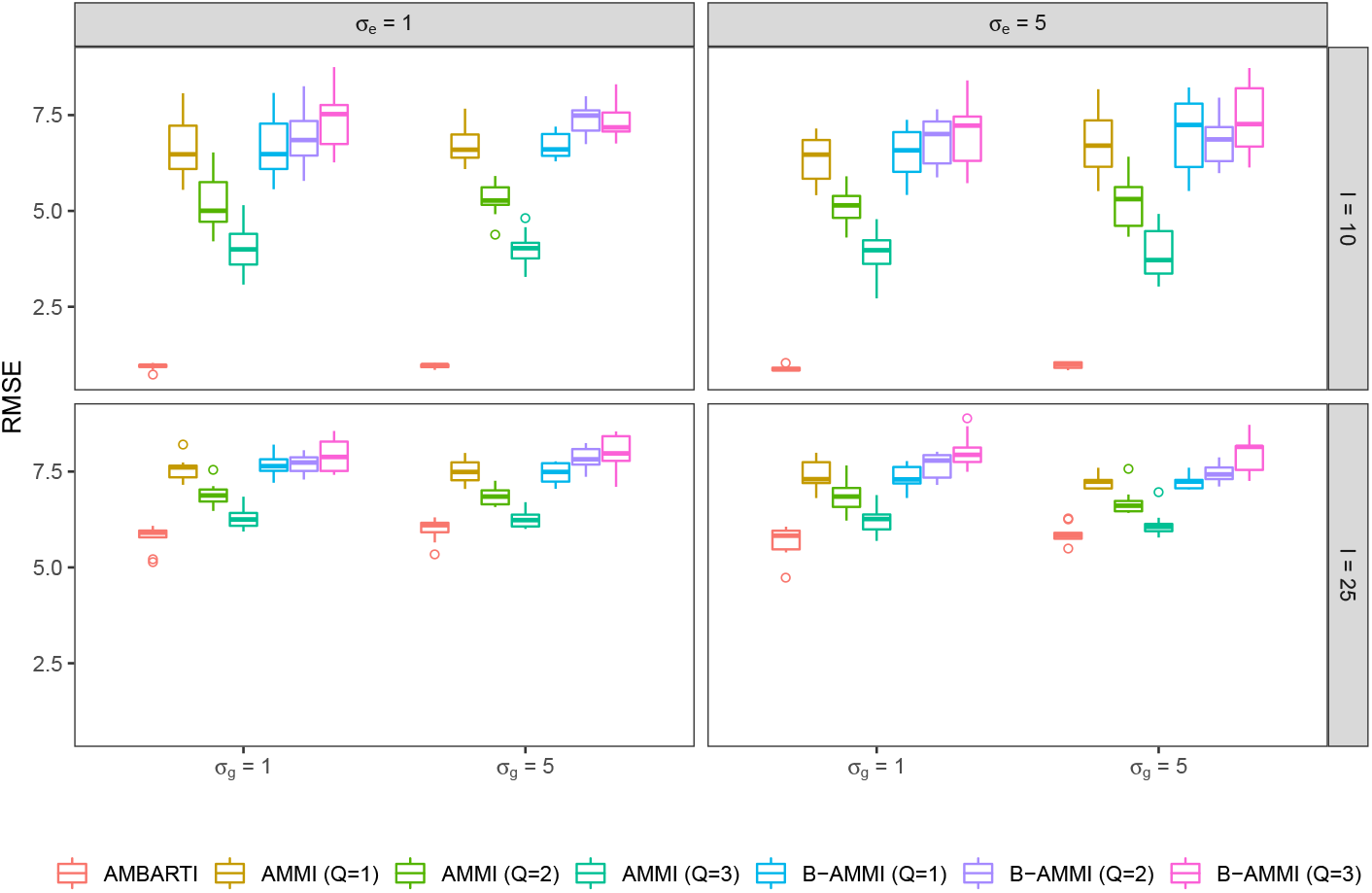
Out-of-sample RMSE related to interaction term of AMBARTI and AMMI models for data simulated from AMBARTI. The different panels show the different parameter values used for the simulation. It appears that the AMMI structure, even with *Q* = 3 cannot capture the interaction behaviour present in the AMBARTI model for *I* = 10.

Figure 5 presents the RMSEs associated solely with the interaction terms from AMMI, B-AMMI, and AMBARTI when the data are simulated from AMMI (which has a bilinear structure for the interactions). The results are presented considering 10 genotypes and 10 environments with different combinations of genotypic and environmental variances. The performance of AMMI is optimal compared to AMBARTI, though the difference between the two is lessened with more complex AMMI model structures (i.e., *Q* = 3). In Figure 6, the values of RMSE are presented for datasets simulated from AMBARTI. In the margins of the figure, the parameters used in the simulations can be found. The RMSE values show that AMMI performs worse than AMBARTI in all scenarios, and in the same cases AMMI RMSEs are three times higher on average than those of AMBARTI for the scenario with *I* = 10 genotypes.

In summary, the information presented in Figures 5 and 6 shows that the AMMI model fails for the complex interactions that can be obtained in the AMBARTI simulated datasets. From a quantitative genetics/biological perspective, there is no reason for the structure of interactions between genotypes and environments to be modelled strictly by a bilinear structure, as more complex structures can be assumed to be present in nature. Thus, AMBARTI may be a more suitable model to estimate the interaction structure in real-world applications.

## 4. Case study: Irish VCU InnoVar wheat data

In addition to the simulation study, real datasets were used to evaluate the performance of AMBARTI. We compare our new approach not only to AMMI and B-AMMI, but also to more sophisticated interaction detection models including smoothing splines ANOVA models (SS-ANOVA; Gu, 2014) and Bayesian multivariate adaptive regression splines (B-MARS; Denison, Mallick and Smith, 1998). To run SS-ANOVA and B-MARS, we used the R packages `gss` (Francom and Sansó, 2020) and `BASS` (Gu, 2014), respectively. A set of value of cultivation and usage (VCU) experiments conducted in Ireland between the years of 2010 and 2019 were considered, and such experiments evaluated the performance of genotypes of wheat *Triticum aestivum L.* across the country for regulatory purposes (i.e., registration of new varieties). Here, our phenotypic response variable is the production of wheat in tonnes per hectare. The design of experiments used was that of a randomised complete block design with 4 replicates. VCUs alongside distinctness, uniformity and stability (DUS) are the most important kind of regulatory multi environmental trials conducted around the world.

The data were kindly provided by the Irish Department of Agriculture, Food, and Marine. Both genotypes and environments were anonymised. These historical Irish VCUs form part of the Horizon2020 EU InnoVar project database (www.h2020innovar.eu). The project aims to build and improve technical solutions for cultivar recommendation based on genomic and phenomic parameters. The models were fitted for all years available (summarised in Table 1), but for brevity we show detailed plots only for the year 2015.

**Table 1.** RMSEfor ŷ on out-of-sample data considering all years in the historical Irish VCU Innovar data. The values of RMSE obtained with AMBARTI are smaller than the ones obtained via AMMI and Bayesian AMMI models (both with Q = 3) for all years considered. The values in bold correspond to the smallest RMSE within each year.

| Year | AMBARTI | AMMI | B-AMMI | SS-ANOVA | B-MARS |
| --- | --- | --- | --- | --- | --- |
| 2010 | **0.81** | 0.83 | 2.21 | 0.82 | 0.87 |
| 2011 | **0.62** | 0.63 | 2.66 | 0.65 | 0.69 |
| 2012 | **0.45** | 0.47 | 2.53 | 0.60 | 0.65 |
| 2013 | 0.57 | 0.57 | 1.48 | **0.55** | 0.56 |
| 2014 | 0.55 | 0.54 | 1.93 | **0.53** | 0.54 |
| 2015 | **0.46** | 0.46 | 1.75 | 0.52 | 0.55 |
| 2016 | **0.44** | 0.45 | 1.98 | 0.50 | 0.50 |
| 2017 | **0.51** | 0.53 | 2.24 | 0.60 | 0.62 |
| 2018 | 0.74 | 0.73 | 1.09 | **0.70** | 0.70 |
| 2019 | **0.56** | 0.58 | 1.26 | 0.72 | 0.70 |

We compare the models by evaluating the estimated values of the genotype and environment effects, and the predictions of interaction behaviour evaluated as: 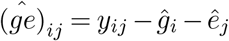. As the Irish data from the InnoVar project correspond to a block design with 4 replicates, to fit all models we randomly selected two replicates and then averaged the response variable across them. This is a common practice in the analysis of GxE experiments with AMMI and B-AMMI models (Josse et al., 2014; Crossa et al., 2011) as these models cannot deal with replicates in an experiment. However, this pre-processing is not needed for AMBARTI. To validate the models, we use the remaining two replicates to calculate the RMSE for *ŷ*.

The estimates of the genotype effects *g_i_* and environment effects *e_j_* are shown in Figure 7 and 8, respectively. The results for AMBARTI show the samples obtained from the posterior distributions, while for AMMI, as it is a frequentist method, we adopted the approach of Goodman and Haberman (1990) where the results correspond to samples from the estimator distributions of *g_i_* and *e_j_*. The rationale of using samples from the estimator distribution is to be able to compare AMBARTI and AMMI results not only in terms of point estimates but also in terms of the uncertainty associated to the point estimates. Although we have tested different number of components *Q* for AMMI, we remark the parameter estimates *g_i_* and *e_j_* do not change regardless of *Q* (i.e., only the bilinear term depends on *Q*; see Equation (2)).

**Fig 7:**
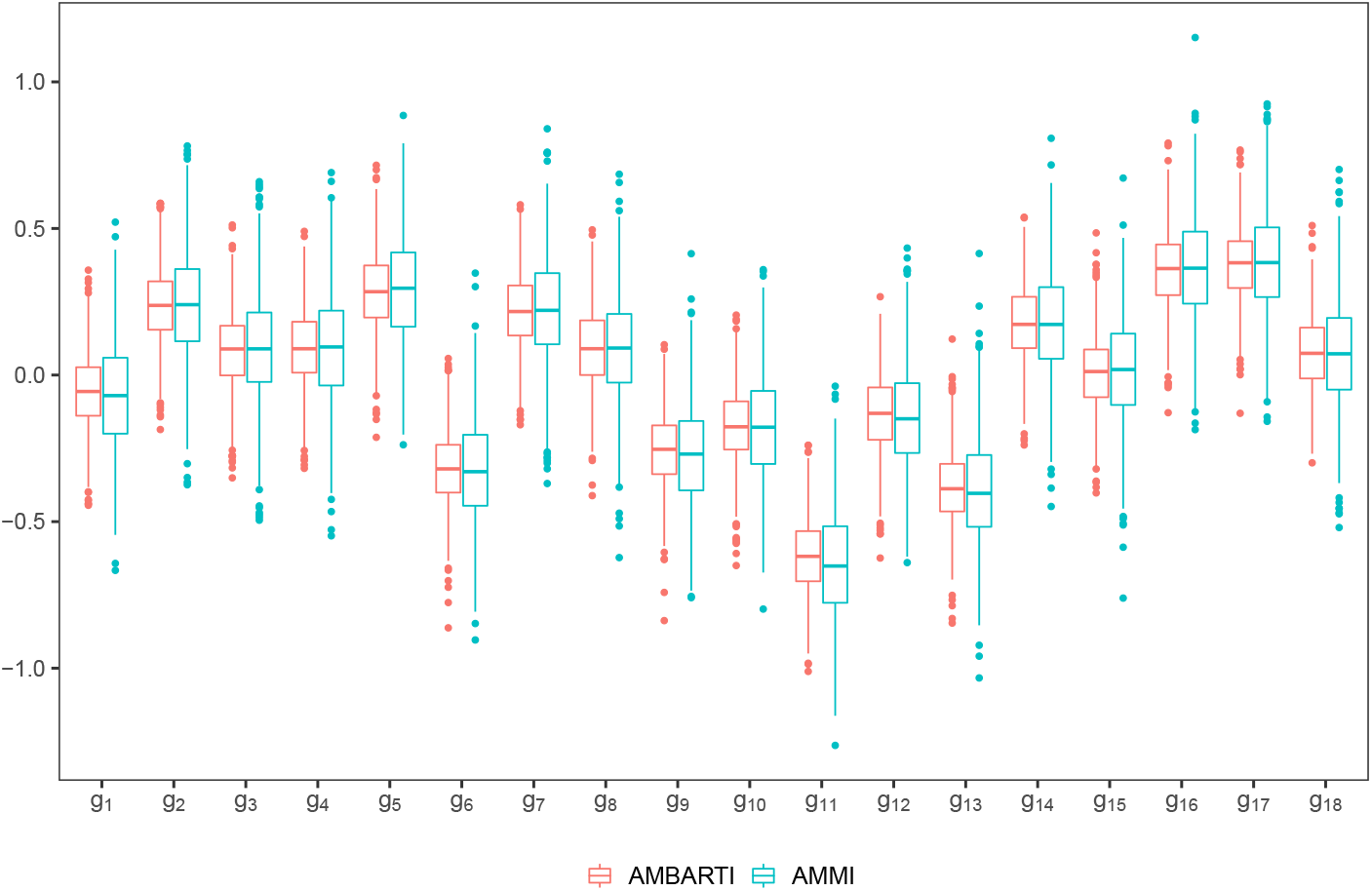
For AMBARTI, the boxplots represent the posterior distribution of genotype effects for the Irish VCU Innovar dataset for 2015. As the parameters estimates from the AMMI model are obtained under the frequentist paradigm, their boxplots summarise samples from the estimator distribution of the genotype effects as presented in Goodman and Haberman (1990).

**Fig 8:**
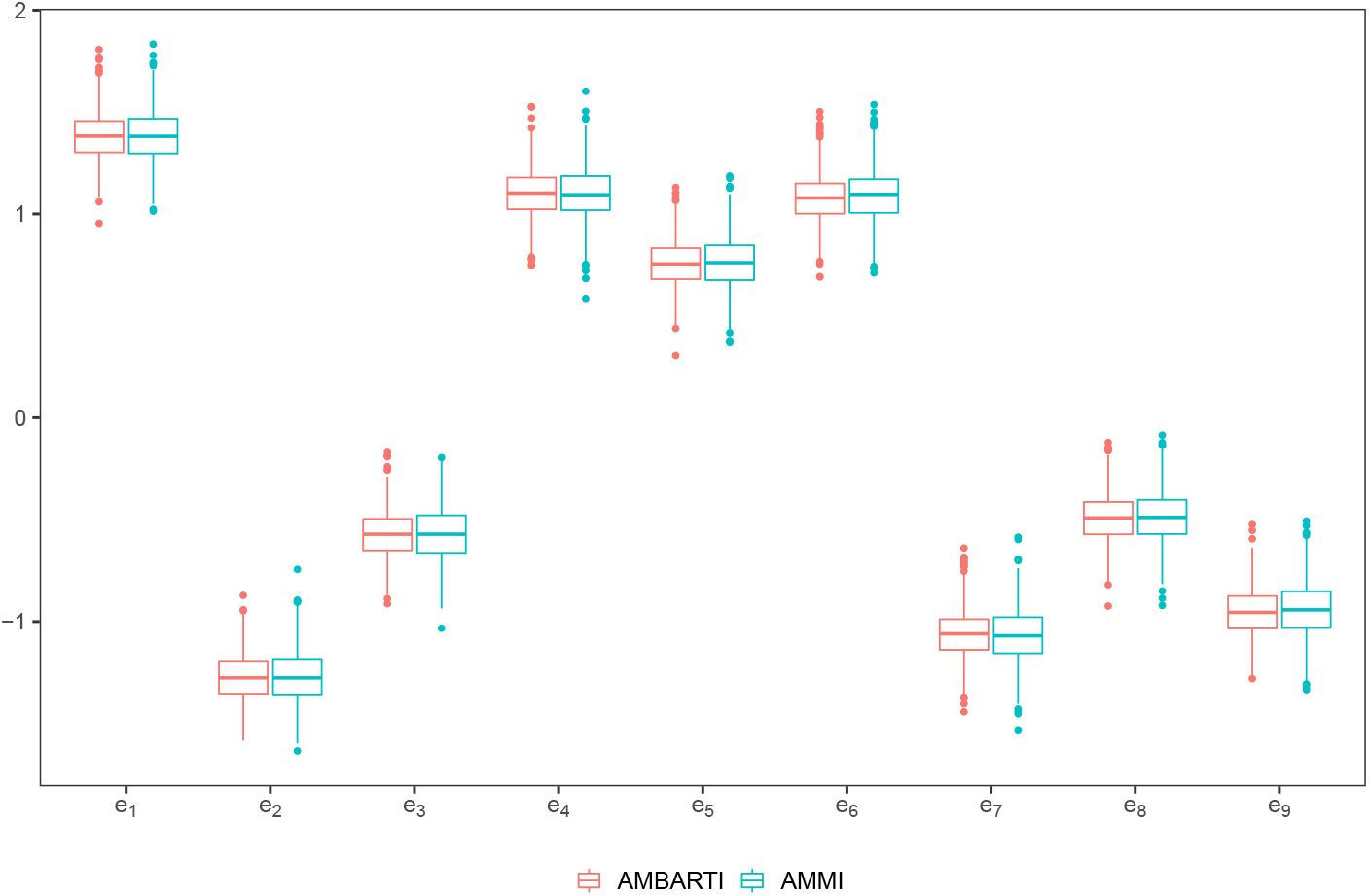
For AMBARTI, the boxplots represent the posterior distribution of environment effects for the Irish VCU Innovar dataset for 2015. As the parameters estimates from the AMMI model are obtained under the frequentist paradigm, their boxplots summarise samples from the estimator distribution of the environment effects as presented in Goodman and Haberman (1990).

In Figure 7, we can see that the credible intervals associated with the main effects of genotypes for AMBARTI are narrower than the confidence intervals from the AMMI model. This is somewhat expected and occurs as BART is able to capture the interactions between *g_i_* and *e_j_* along the MCMC process, which in turn decreases the residual variance *σ*^2^. Hence, as the full conditionals of the *g_i_* and *e_j_* depend on *σ*^2^, their estimates become less uncertain as *σ*^2^ gets smaller; see the supplementary material (Sarti et al., 2022) for details on the AMBARTI algorithm. For the sake of visualisation, we decided to omit the B-AMMI results as they presented too many values that exceeded —10 and 10.

A more complete comparison across all years is shown in Table 1. In this table, we calculated the predicted values *ŷ* on the out-of-sample data. We can see that RMSEs obtained with AMBARTI are smaller than the ones returned by the AMMI model for all years, thus highlighting that the AMBARTI model can more accurately estimate the marginal effects along with interaction component. Further, we can note that the B-AMMI presents the worst results among all methods considered. One could expect the results from AMMI and B-AMMI would be similar, but the Bayesian version is different in spirit to its frequentist counterpart, as it does not estimate the main effects and bilinear term in a two-stage approach. Instead, B-AMMI obtains the parameters through a one-stage procedure and assumes priors that do not take into account the orthormality constraints, which are fundamental for the identifiability of the frequentist AMMI model. In the B-AMMI, the orthormality constraints are applied via a post-processing on the parameter estimates after the model is fitted. However, we observed the B-AMMI model with and without the post-processing may have very different performance.

Although the results for SS-ANOVA and B-MARS are competitive with AMBARTI, they work differently and do not return results in the format needed to fit the purpose of the analysis. For instance, SS-ANOVA returned interactions that do not make sense (e.g., interactions between environments or interactions between genotypes). In contrast, B-MARS is similar to BART in the sense that it does not require any specification but with the advantage that it decomposes the impact of individual and interaction effects on the response.

Regarding the computational time, AMBARTI took about 6 minutes on average to run, considering 50 trees, 1000 iterations as burn-in and 1000 iterations as post burn-in. This time was registered in a MacBook Pro 2.3 GHz Dual-Core Intel Core i5 with 8GB memory. AMMI took just seconds. This difference could be reduced by optimising the AMBARTI implementation using routines in C++ similar to those for BART implementations in R packages `BART` (McCulloch et al., 2020) and `dbarts` (Dorie, 2020). However, we believe AMBARTI’s superior performance and posterior estimation of uncertainties outweighs the longer computational time.

### 4.1 New visualisations for AMBARTI main effects and interactions

One of the key outputs of the standard AMMI model is the biplot (Gabriel, 1971), which assists in the determination of important GxE interactions and may be used for cultivar recommendation. However, these plots display only the interaction measure, thus missing the key marginal effects that may also come into play. For example, a certain genotype and environment may have a strong positive interaction, but if the genotype is consistently poor in all environments this may not be clear in the biplot. Instead, we introduce new types of plots that give this full consideration.

Our first new plot is based on a heat map adapted to display both the GxE interactions (along with the marginal effects) and the predicted yields from the AMBARTI model. In Figure 9, we display the GxE interactions in the centre of the plot and the marginal effects for both environment and genotype as separate bars in the margins. The ordering applied to Figure 9 is in terms of the marginal effects for both environment and genotype, and displays low values in red to high values in blue. As both the GxE interactions and marginal effects are on the same scale and are centred around zero, we display them using only one legend and use a divergent colour palette. A diverging colour palette uses two diverging hues to represent the extremes and highlights the midpoint with a light colour. This allows for quick identification of the GxE interactions and to observe which of the environments or genotypes are the most or least optimal.

**Fig 9:**
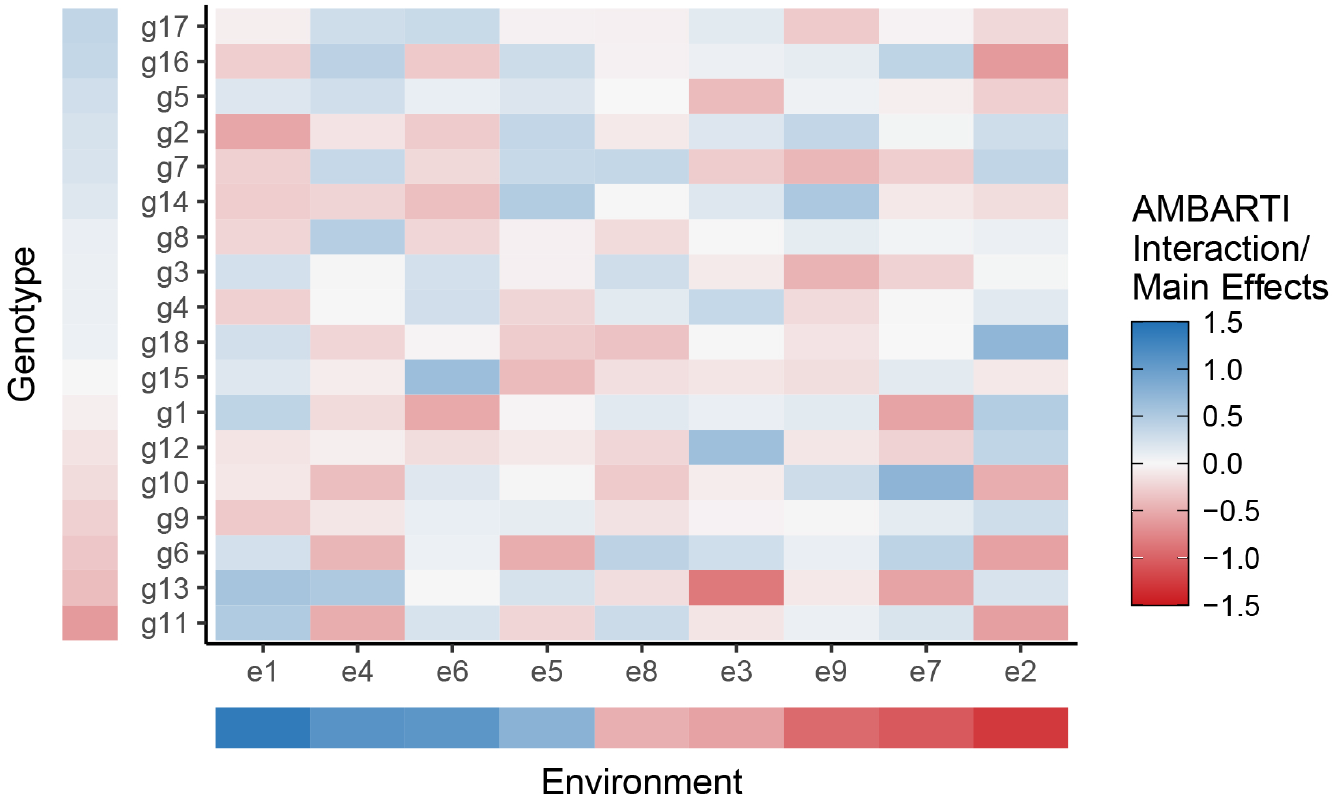
GxE interactions and main effects for the AMBARTI model sorted by the main effects for the Irish VCU InnoVar data in 2015. We can clearly see that environments 1, 4, 6, and 5 provide superior yields for many of the genotypes studied. Furthermore, environment 1, for example, seems to interact particularly strongly in a negative way with genotype 2. The grand mean μ is not included in this plot for ease of identification of marginal and interacting effects.

In Figure 10, we show the ordered heat map of the predicted yields (as opposed to their component parts shown in Figure 9) for each combination of environment and genotype for the AMBARTI model. In this case, we use the same ordering as that in Figure 9 with high values being generally displayed in the top left, moving to low values at the bottom right, with the units for the plot being the same as that of the phenotype (i.e., yield/production of grains in tonnes per hectare). For this plot, we use the same diverging colour palette as in Figure 9 as the scale is centred around the mean, and when combined with the ordering, this gives a clear identification as to which environment and genotype produce high or low yields. Additionally, the AMBARTI methodology makes it possible to use tree methods to identify which combinations of genotypes and environments are similar considering a given phenotypic characteristic, e.g., yield. For example, the combination of genotype 1 and environment 1 and genotype 17 and environment 1 are similar in terms of predicted yield, as shown by the colours in Figure 10. Similarly, genotype 2 has similar yields for environments 4 and 5. Thus, we can consider these environments similar and group them in a set named mega-environment 4-5 considering genotype 2. Such a concept is crucial for cultivar recommendations. To visualise the uncertainty associated with Figures 9 and 10, we provide plots which show the median, 5%, and 95% quantiles for the predicted response and GxE interactions and main effects in the supplementary material (Sarti et al., 2022).

**Fig 10:**
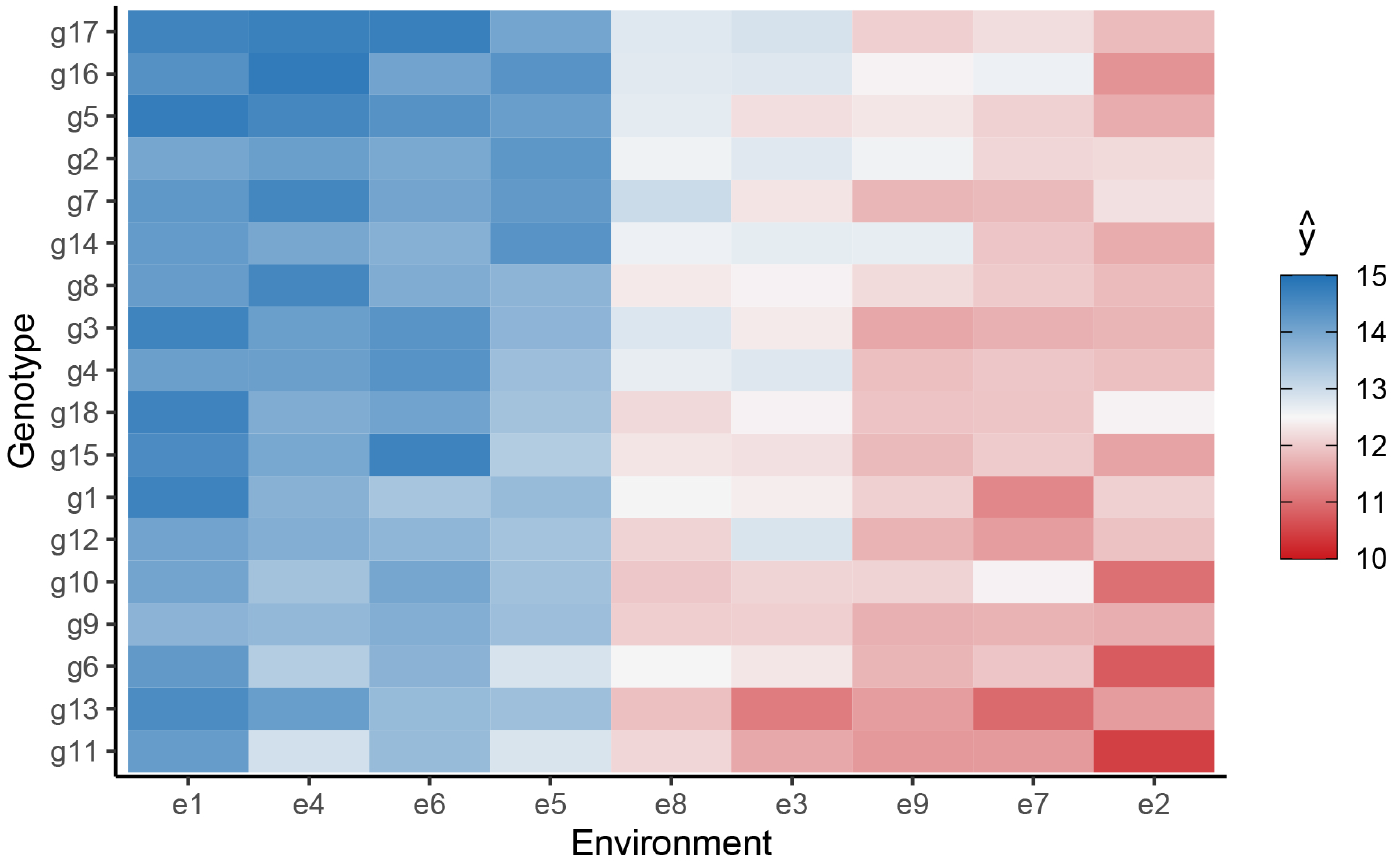
Predicted yields from the AMBARTI model for the Irish VCU InnoVar data in 2015. Values are sorted by the main effects. We can see, for example, a high value for the predicted yield for environment 1 with genotype 5 and a low value between environment 2 and genotype 11.

In Figure 11, we show a bipartite plot of the information displayed in Figure 9, but showing only the extremes of the high and low values. In this case, we display just the top 2% and the lowest 2% of the interactions. We employ the same diverging colour palette as Figure 9 except in this case the colour of the nodes represent the marginal effects and the size of each node represents the absolute value of the marginal effects. Similarly, the colour of the connecting edges represent the interaction values and the width of each edge represents the absolute interaction value. That is, larger magnitudes of the marginal effects will result in larger nodes (and vice-versa), and larger magnitudes of the interactions will result in thicker edges (and vice-versa). The aim of this plot is to allow the reader to easily and quickly identify which of the environments are the most and least optimal for each genotype and to also identify where there are clear interactions. Quantile versions of Figure 11 could also be plotted to assess uncertainty.

**Fig 11:**
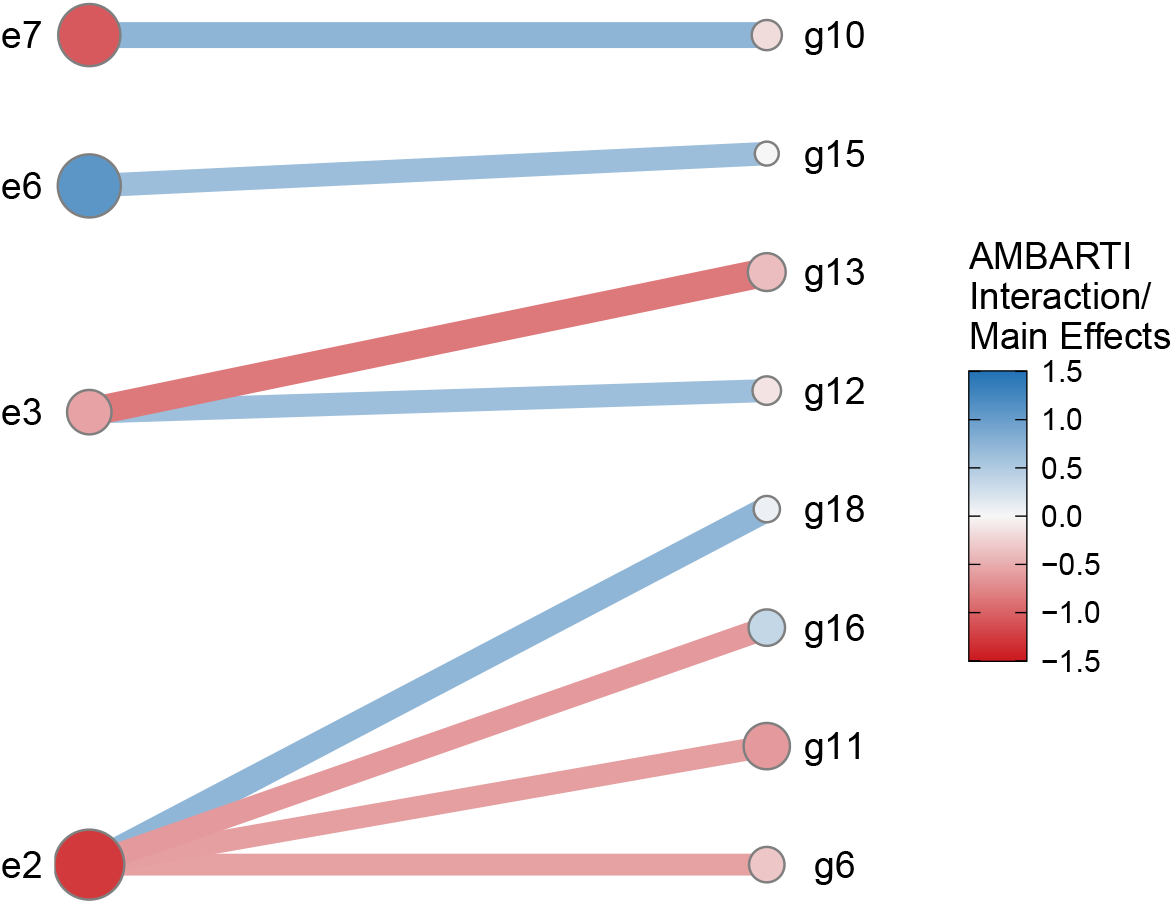
Bipartite network plot showing the top (in blue) and bottom (in red) 2% GxE interactions and main effects from the AMBARTI model for the Irish VCU InnoVar data in 2015.. We can see that environment 3 has strong positive and negative interactions with genotypes 12 and 13, respectively.

The visualisation perspective proposed here helps construct easily interpretable agronomic recommendations. Figure 10 can help users with no background in statistics identify that the best genotypes considering yield are the ones in the top left corner: *g*_17_,*g*_16_,*g*_5_,*g*_2_. These genotypes will have a tendency to have a better acceptance by farmers, considering solely the yield in tonnes per hectare assuming higher yields are economically preferred. Figure 9 shows us that environments *e*_1_, *e*_4_, *e*_6_, *e*_5_ are related to higher marginal effects and should be considered preferential to crop the list of wheat genotypes evaluated.

Figures 9 and 11 are also useful to establish combinations of genotypes and environments that should be avoided when the interaction is negative, indicating that a given genotype does not perform well in a given environment. This negative interaction increases the risk of low yield and consequent economic impacts. Combinations to be avoided exist even for environments and genotypes with high marginal effects. For instance, the combination of *g*_2_, *e*_1_ should be avoided even though *g*_2_ and *e*_1_ have high marginal effects. This is an important information for regulators who may be responsible for a variety’s commercialisation approval or agents that promote credit or insurance for farmers given to risks that the negative interaction implies. Farmers who produce a genotype not indicated for their environment can end having a worse score or risk. On the other hand, Figure 10 is also useful to spot the combinations of genotypes and environments that should be encouraged once the signal of the interactions is positive.

In adaptability breeding, the breeder seeks to find the best genotype for a specific environment or a small set of environments. In broad target strategies, the aim is to find genotypes that perform well across several environments. For example, in Figure 9, *g*_5_ has high marginal effect and performs well (and interacts positively) with environments *e*_1_, *e*_4_, *e*_6_, *e*_9_. Similarly *g*_16_, the second best genotype considering marginal effects, performs well in environments *e*_4_, *e*_5_,*e*_9_,*e*_7_. Genotypes which present better performance across several environments are classified as high stability genotypes. They tend to be preferred by breeders because they allow optimisation of processes in the chain of seed production.

## 5. Discussion

We have introduced a new model named additive main effects Bayesian additive regression trees interaction (AMBARTI). AMBARTI is a cut Bayesian semiparametric machine learning approach that estimates main effects of genotypes and environments and interactions with an adapted regression tree-like structure. This approach to interactions allows the treatment of more complex structures than the ones considered by traditional models.

Given the fact that GxE interactions are the result of a tangled myriad of genetics, proteomics, biochemical, environmental and additional factors, the flexibility of AMBARTI in dealing with more complex interactions can be seen as an important improvement in the understanding of the complexities associated to GxE phenomenon. In practice, the choice between a low-rank model (AMMI) and a model that is able to deal with a sparse interaction structure (AMBARTI) is a modelling choice. However, in the real data examples upon we have tested the models, AMBARTI performed slightly or much better than AMMI, which suggests that a bilinear term is perhaps an oversimplification to study the relations that arise from GxE interactions. We believe that AMBARTI is a useful candidate to expand the understanding of experimental data in quantitative genetics.

The main novelty in AMBARTI comes from its semi-parametric structure which enables the uncertainty to be shared between the main effects and the interaction trees. More specifically, we design the trees so that they can only incorporate the interaction terms by forcing each branch of a tree to split on both a genotype and an environment. We have shown in simulation experiments that this yields similar estimates to traditional models for the marginal effects and superior estimates for the interaction terms, which are no longer restricted to be linear in a restricted dimensional space. This removes the need for, e.g., the arbitrary selection of the *Q* parameter in a standard AMMI formulation.

A second novelty is that we have introduced new displays that simultaneously allow for interpretation of the marginal and joint effects. We have created both a heatmap and a bipartite network-style plot of the results, which are not limited for use with an AMBARTI model and could be applied to any suitable model (for example, AMMI, B-AMMI, etc.) and are a useful tool for deciphering complex model structures. From these plots, we hope to enable those using the output of AMBARTI models to make more informative decisions about which genotype and environments are most compatible.

We believe that there are many possible extensions of the AMBARTI approach. Other more advanced methods, such as PARAFAC (Basford, Kroonenberg and DeLacy, 1991; Harshman and Lundy, 1994), are available for higher dimensional interactions (such as with time). These are different versions of tensor regression (Guhaniyogi, Qamar and Dunson, 2017) and, in theory, there is no reason why the AMBARTI approach cannot be used for higher dimensional tensor-type interactions, though this is not currently possible in our code. Similar enhancements for multivariate outputs and time-series like-structure seem promising, and we hope to explore these in future work. As suggested by a referee, modelling approaches which do not rely upon the BART model could also be explored. For instance, the spike-and-slab priors (George and McCulloch, 1997; Ishwaran and Rao, 2005) commonly used for Bayesian variable selection could be adapted to identify important GxE interactions. Based on some preliminary analyses, the results were promising and this a subject of ongoing work.

## Acknowledgments

Danilo and Estevão are joint first authors. We would like to thank the editor and the two anonymous referees for their thorough comments, suggestions and ideas that greatly improved the manuscript. We are very grateful to John Joe Byrne at the Department of Agriculture for providing us with the dataset for our case study. Danilo Sarti and Antônia A. L. dos Santos received funding for their work from the European Union’s Horizon 2020 research and innovation programme under grant agreement No 818144. Alan Inglis and Estevão Prado’s work was supported by the Science Foundation Ireland Career Development Award grant number 17/CDA/4695 and SFI research centre 12/RC/2289_P2. Andrew Parnell’s work was supported by: the Science Foundation Ireland Career Development Award (17/CDA/4695); an investigator award (16/IA/4520); the Marine Research Programme funded by the Irish Government, co-financed by the European Regional Development Fund (Grant-Aid Agreement No. PBA/CC/18/01); European Union’s Horizon 2020 research and innovation programme under grant agreement No 818144; SFI Centre for Research Training 18CRT/6049, and SFI Research Centre awards 16/RC/3872 and 12/RC/2289_P2. For the purpose of Open Access, the author has applied a CC BY public copyright licence to any Author Accepted Manuscript version arising from this submission’.

## SUPPLEMENTARY MATERIAL

In the supplementary material (Sarti et al., 2022), we provide details about the AMBARTI implementation (Section A), the naive and tempered cut algorithms (Section B), orthonomality constraints in AMMI models (Section C) and visualisation of the uncertainties of the parameter estimates in the AMBARTI model (Section D).

## SUPPLEMENTARY MATERIAL

### 1. AMBARTI implementation

In this Section, we detail the AMBARTI model. Firstly, the conditional probability distribution associated with *y_ij_* is

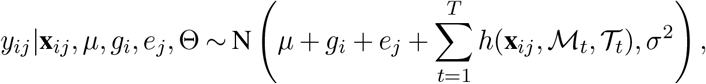

where *y_ij_* denotes the response for genotype *i* and environment *j*, 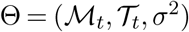, *μ* is the grand mean, **x**_*ij*_ is the row of the design matrix **X** associated to observations with genotype *i* and environment *j*, and *h*(·) = *μ_tℓ_* is a function that assigns the predicted values 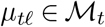 to observations that belong to 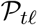, with 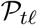 denoting the set of rules that define the node *ℓ* of the tree *t*. In order to obtain the posterior distributions needed for the model, we assume the following prior distributions:

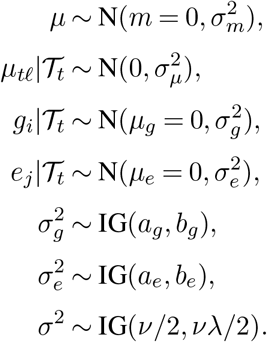

The prior distribution on the tree structure depends on the depth and number of terminal and internal nodes, and is given by

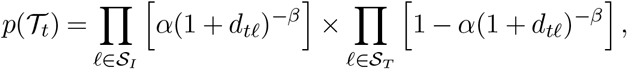

where 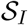 and 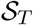 denote the sets of indices of the internal and terminal nodes, respectively, and *d_tℓ_* represents the depth of the node *ℓ* of the tree t. Furthermore, let 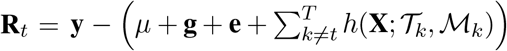 denote the vector of the partial residuals, where **g** and **e** are vectors containing the main effects *g_i_* and **e*_j_* for all *i* and *j*. Below, we present the cut full conditional of *μ*.

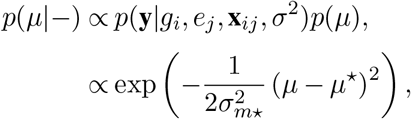

which is a

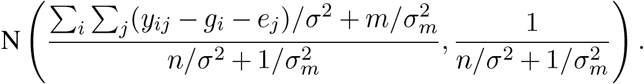

Hence, the cut full conditional of *g_i_* is given by

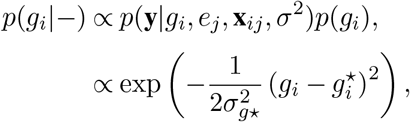

which is a

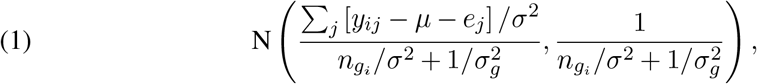

where *n_g_i__* is the number of observations that belong to *g_i_*; similarly to *n_*e*_j__*. Analogously, the cut full conditional of **e*_j_* can be written as

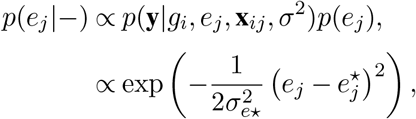

which is a

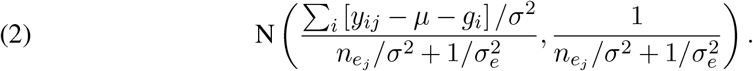

The full conditional of 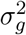 is given by

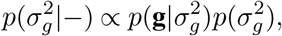

which is an

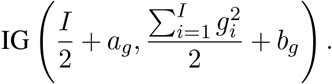

The full conditional of 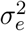 is written as

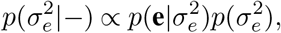

which is an

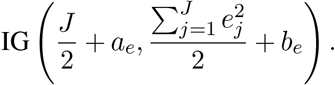

In addition, we present the full conditional of the trees. This distribution is utilised to compare the previous tree to the current one, as in BART the splitting rules are created by randomly selecting a covariate and a split point. Below, we present the full conditional of 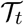 as

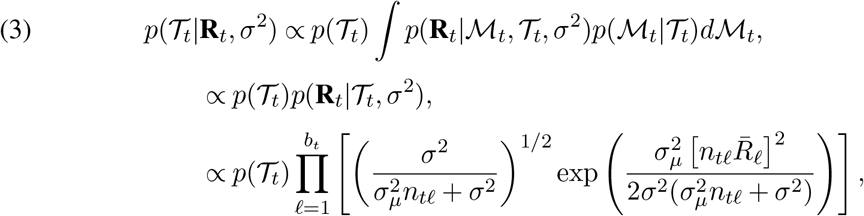

where 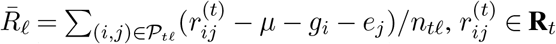 and *n_tℓ_* is the number of observations that belong to 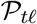. To sample from this expression, the Metropolis-Hastings algorithm is used, because a closed-form distribution is not obtained in this case.

As all *μ_tℓ_* are i.i.d, it is possible to write 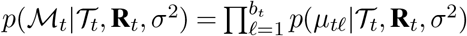. Similarly to the original BART, the full conditional of *μ_tℓ_* in the AMBARTI model also depends only on the information provided by all trees, except by 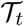, via partial residual as **R**_*t*_. Hence, the full conditional of *μ_tℓ_* can be written as

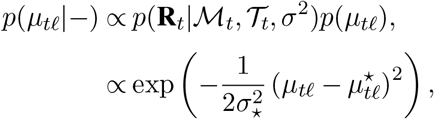

which is a

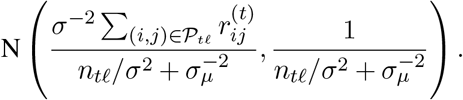

Finally, after generating all predicted values for all trees, *σ*^2^ can be updated based on

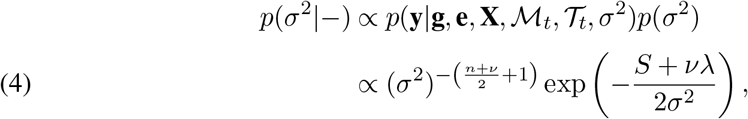

where 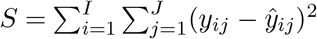 and 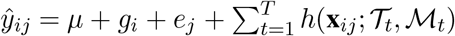. The expression in (4) is an IG((*n* + *v*)/2, (*S* + *νλ*)/2). In algorithm 1, the AMBARTI model is presented for one MCMC iteration.

#### Algorithm 1: AMBARTI Algorithm

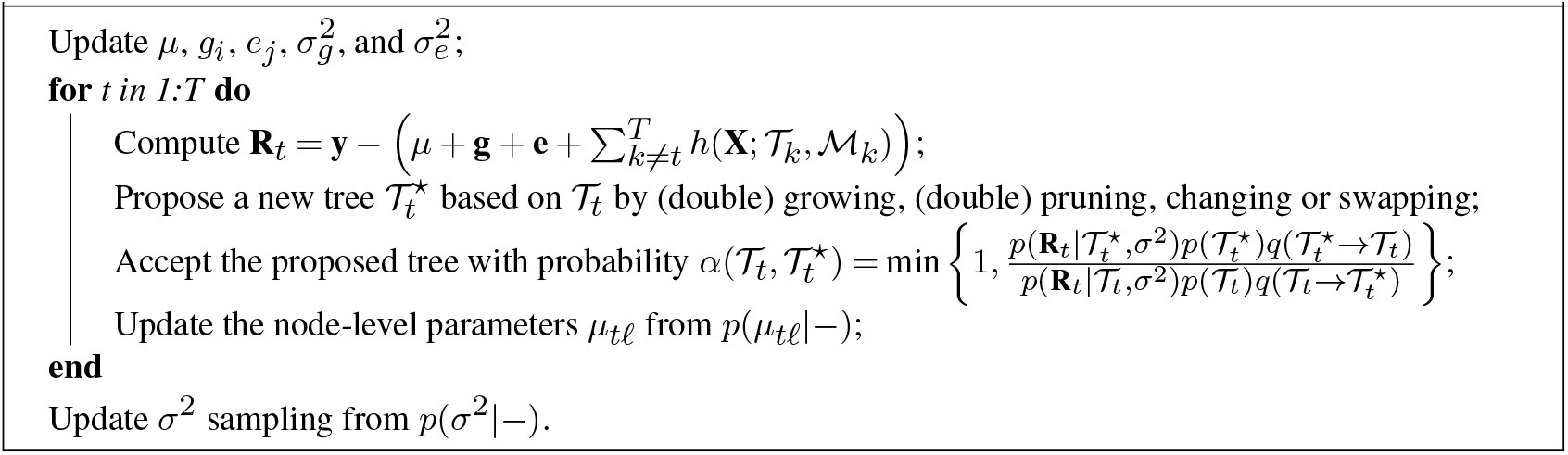

### 2. Naive and tempered cut algorithms

We performed a simulation experiment to compare the results from the naive and tempered cut algorithms. We did this because the results in the paper are all based on the former method. The rationale of the tempered transitions, which are obtained from the tempered cut algorithm, is instead of moving from, say, 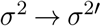, we move along a linear path in a sequence 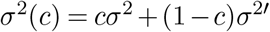, where *c* = (*c*_1_,…, *c_m_*) with *c_k_* = *k*/*m*. Then, at each step, conditioned on *σ*^2^(*c*), a sample for 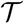 is obtained but only the last one is kept, which becomes 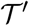 (Plummer, 2015). In this way, the condition in (9) in the main text is satisfied and approximate samples from the cut posterior distribution of interest are obtained. Since 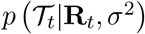 in AMBARTI does not have a known distributional form and due to the use of a cut model to avoid the feedback between the main effects and the BART component, Plummer (2015) recommend performing tempered transitions for *g_i_*, **e*_j_* and *σ*^2^.

We have carried out this comparison to show whether the posterior samples from 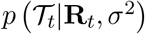 differ under each method. Recall that in AMBARTI the following parameters have closed-form full conditionals: *g_i_* (effect of genotype), **e*_j_* (effect of environment), *μ_tℓ_* (BART predictions), *σ*^2^ (residual variance). Thus, to sample from 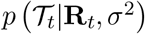 via the tempered cut algorithm, the transitions from 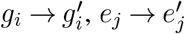 and 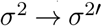 need to be smooth and cannot lead to large jumps. We point out that there is no need to adjust the transitions from 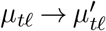 since these are marginalised out in 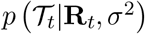 as shown in (3).

Unlike the other full conditionals in the AMBARTI model, the full conditional for the trees is used as a mechanism to filter ‘good’ splitting rules only, as opposed to returning a value from the corresponding posterior distribution. Thus, to examine the convergence of the trees is not a straightforward task because i) the splitting rules are randomly selected and, as a consequence, ii) there is not a conventional MCMC chain of posterior samples to be examined. To illustrate this point, suppose that we ran AMBARTI more than once and then examined the structure of, say, 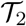. The structure/splitting rules of 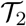 certainly will not be the same across the runs, especially if the number of covariates is large. This is not necessarily an issue and happens frequently due to the uniform specification on the splitting rules where independent uniform priors are placed on the available covariates and split values. Conversely, the convergence of the posterior samples of *g_i_*, **e*_j_* and *σ*^2^ can be assessed by visualising the posterior density of these parameters along the line of Figure 4 of Plummer (2015).

Figure 1 in the main text shows some elements of the tree structure that could be used in an attempt to evaluate the convergence of the lives. Once again we point out that the predicted values *μ_tℓ_* are integrated out from 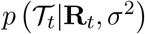. The remaining quantities, which are derived from the tree structure rather than parameters directly monitored during the MCMC run, are the depth of the tree, the number of terminal nodes (circles), the number of internal nodes (rectangles) and the splitting rules, which are represented into the internal nodes.

The maximum depth of a binary tree is directly related to the number of nodes in the tree. In addition, the number of internal nodes in the same type of tree is equal to the number of terminal nodes minus 1. Thus, in an attempt to evaluate the convergence of the trees, it seems reasonable to look at the distribution of the number of terminal nodes, based on the aforementioned relations. More specifically, we are interested in empirically evaluating whether the distribution of the number of terminal nodes changes as the number of steps in the tempered transition increases for the tempered cut algorithm.

With this in mind, we simulated synthetic data from the AMMI model as it is the least favourable scenario for the AMBARTI model due to the orthonormality constraints on the bilinear term. The data were simulated considering *I* = 10 genotypes, *J* = 10 environments, *Q* = 3, λ = (12,10,8), and standard errors 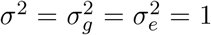. Furthermore, we have analysed the distribution of the number of terminal nodes for our case study Innovar data set for 2015. For both synthetic and real data, we used an increasing number of steps (*m* = 1,2,4,10,20, and 40). We have considered steps greater than 40, but the results were not any different compared to 40.

Figure 1 shows the results of the mean number of terminal nodes for the different number of steps *m* in the tempered cut algorithm. In panel (a), the results are for the simulated data, while in panel (b) they correspond to the Innovar data. As we can see, the distribution of the number of terminal nodes in both panels do not differ between *m* = 1 (naive cut algorithm) and the other number of steps (tempered cut algorithm). For instance, in panel (a), we can see that the boxplots for *m* = 1,20 and 40 are similar in terms of the first, second and third quartiles, which highlights that the tempered transitions have no effect on the topology of the trees in AMBARTI for these data. In panel (b), the difference between the results of *m* = 1 and the others number of steps is negligible. Finally, we have also noticed that the tempered transitions (with steps greater than 1) have not changed either the estimates of the additive effects, the estimates of the interactions, the overall fitted values, or even the posterior prediction intervals.

**Fig 1:**
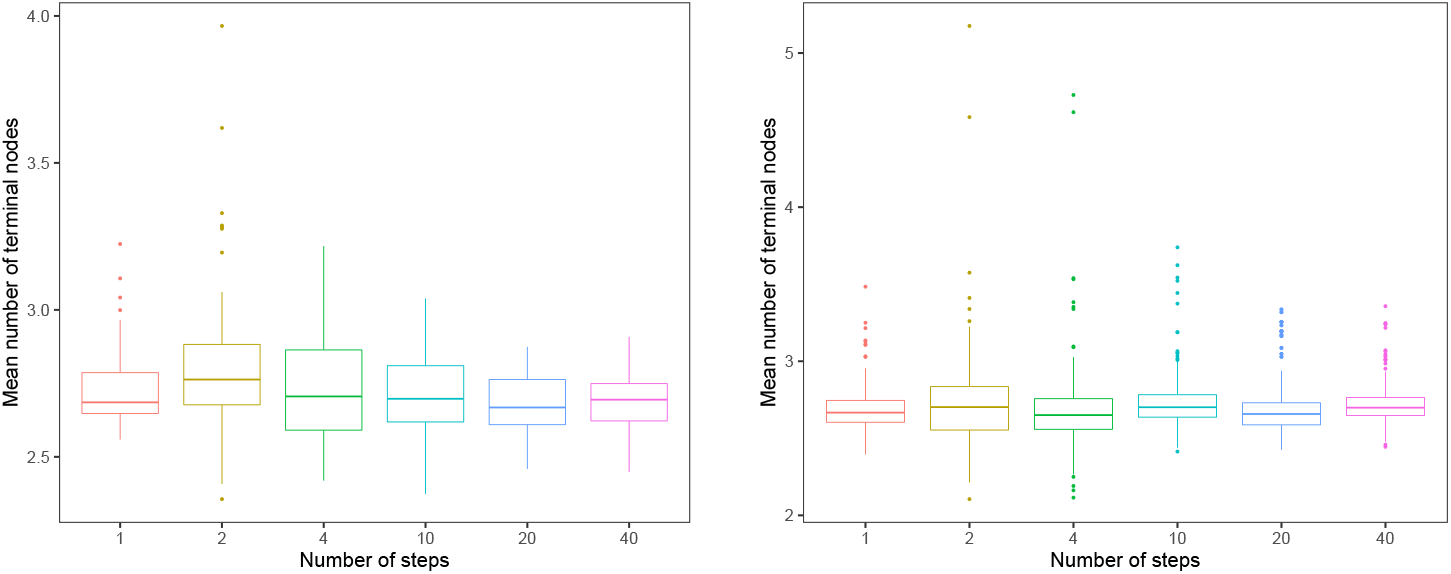
Mean number of terminal nodes in the AMBARTI model with tempered transitions. The numbers of steps used in the tempered cut algorithm are displayed on the x-axis. The y-axis shows the mean number of terminal nodes observed after the burn-in period. Each boxplot contains 200 values corresponding to the mean number of terminal nodes in each tree used in the ensemble of the BART component. In panel (a), the results are shown for the synthetic data simulated from the AMMI equation. In panel (b), the results are for the Innovar data for 2015.

Given the similarity of the distributions of the number of terminal nodes under the naive and tempered cut algorithms for the data considered above, we kept the results from the naive cut algorithm in the simulation and case study sections. In practice, the results shown in this Section provide elements that indicate that the condition in (9) in the main text, after an appropriate burn-in period, seems reasonably attained even under the naive cut algorithm for the AMBARTI model.

We believe the explanation for the similar results is two-fold. First, the transition 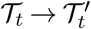 for 50% of the moves (i.e., change and swap) in the BART component have no dependence on the previous tree 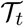 since the ratio of the transition kernel for these moves is 1. Second, the prior on the trees forces them to be shallow and this is reflected in the low mean number of terminal nodes per tree. Unlike the cervical cancer example of Plummer (2015) where the Metropolis algorithm is set up to give acceptance probabilities in a certain range for the (cut) parameter of interest, in AMBARTI it is not possible to control the acceptance probability rate of the Metropolis step because the splitting rules are randomly proposed.

Although the tempered/naive cut algorithms produced similar results, we point out that the number of steps in the tempered transitions needs to be assessed for the application at hand. In our software, we have added an argument called `nsteps` which allows the user to manually specify the number of steps in the tempered transitions, with the default option being `nsteps=1`.

### 3. Orthonormality constraints of the AMMI model

We recall the AMMI model is overparameterised, so constraints need to be imposed so that the parameters can be estimated (Josse et al., 2014). In this Section, we show how to apply the orthonormality constraints on *γ_iq_* and *δ_jq_* when simulating from the AMMI model.

Let *γ* be an *I* × *Q* matrix, ***δ*** a *J* × *Q* matrix, and consider that *γ_iq_* and *δ_jq_* are elements in row *i* and column *q* of the corresponding matrices. The following constraints are considered: i) 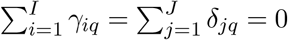, for *q* = 1,…,*Q*; ii) 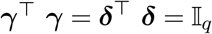, where 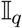 represents an identity matrix of dimension *q*; iii) *λ*_1_ ≥ λ_2_ ≥ … *λ*_*q*-1_ ≥ *λ_q_* ≥ 0 and; iv) *γ*_1*q*_ ≥ 0, for all *q* = 1,…,*Q*.

To illustrate our the strategy to meet the constraints presented above, we take the *γ_iq_* as an example, but this also works for *δ_jq_*. First, we create *S* an *I* × *Q* matrix, where 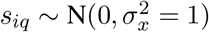. Here, *s_iq_* could be sampled from other distributions, such as Gamma or Beta. In addition, we define 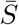 as an *I* × *Q* matrix with each element being the mean of the corresponding *q* column of *S*. Hence, we know that

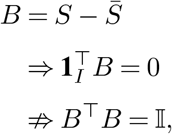

where **1**_*I*_ is a column vector of dimension I containing ones, *B* is, by construction, a full rank matrix and *B^T^B* is symmetric. However, we find a matrix *A* such that 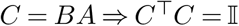. That is, we know that

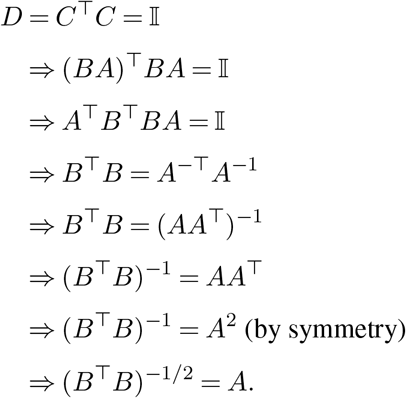

In the end, we have that *γ* = *B*(*B^T^B*)^1/2^.

### 4. Visualising the uncertainties of the parameter estimates in the AMBARTI model

**Fig 2:**
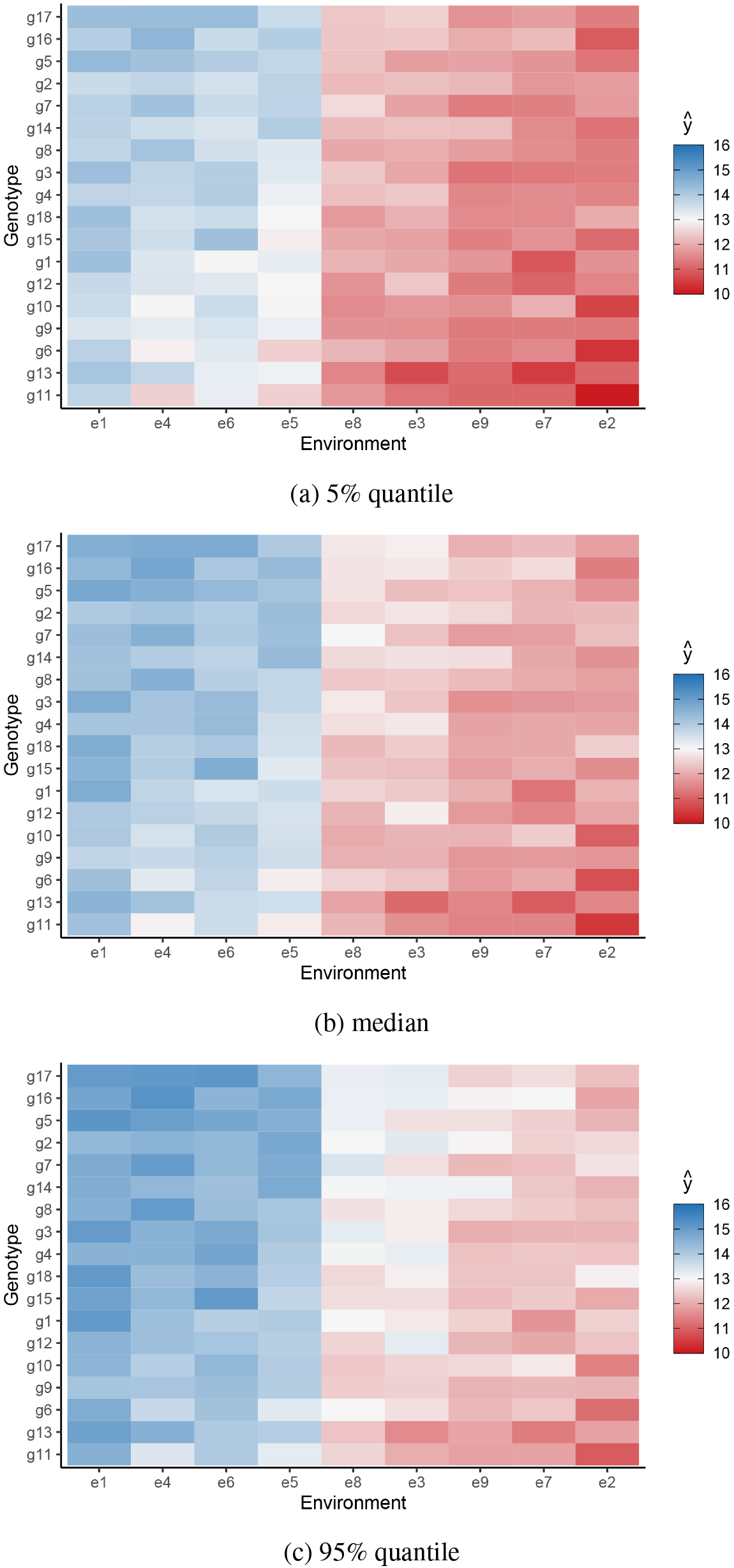
5%, median, and 95% quantiles for predicted yields from the AMBARTI model for the Irish VCU InnoVar data in 2015. The three parts of the graph allow us to address the uncertainties associated with the predicted response in yields described in Figure 10 in the main text.

**Fig 3:**
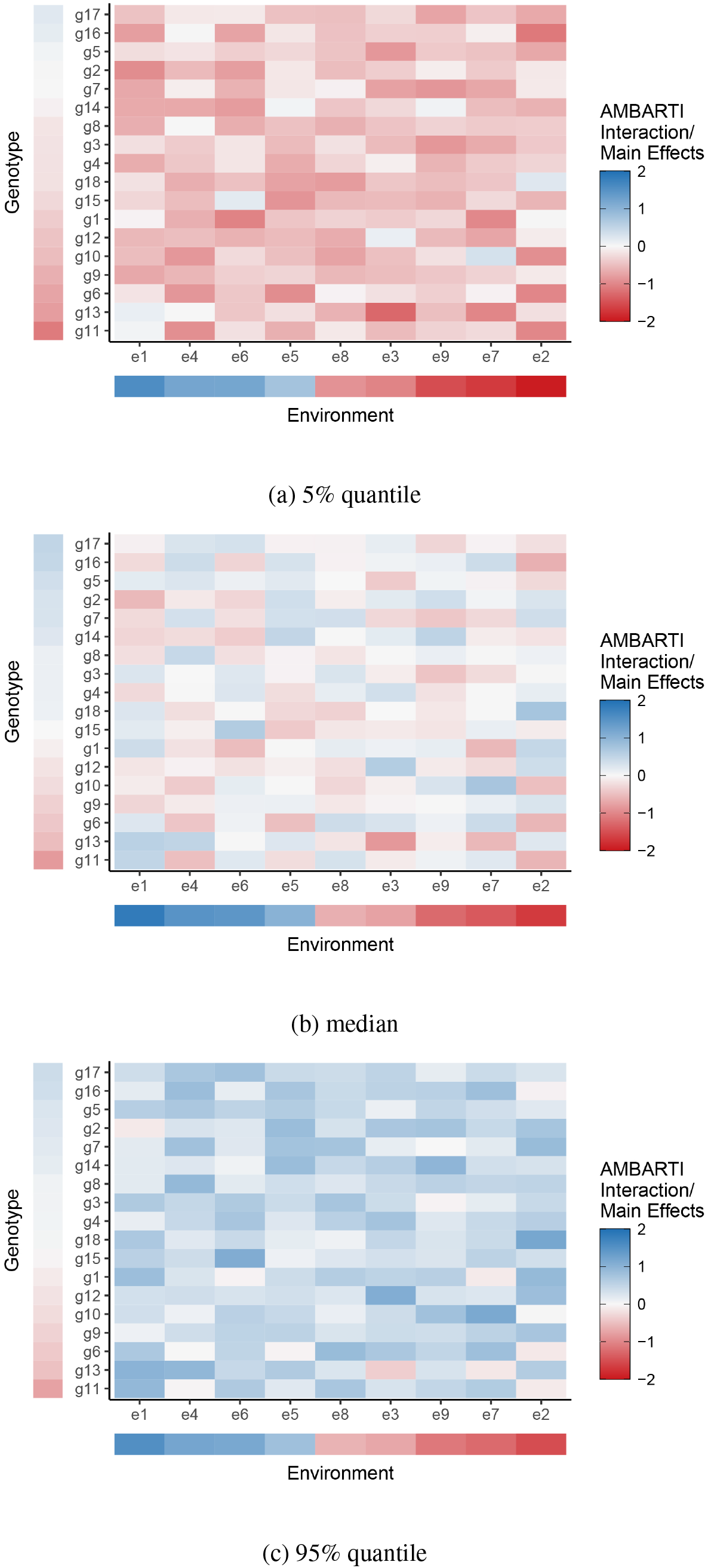
5%, median, and 95% quantiles for predicted yields from the AMBARTI model for the Irish VCU InnoVar data in 2015. The three parts of the graph allow us to address the uncertainties associated with the GxE interactions and the main effects obtained by AMBARTI. We can see that the credible interval associated to *g*_18_/*e*_2_, *g*_10_/*e*_7_, *g*_13_/*e*_1_, *g*_11_/*e*_1_, and *g*_13_/*e*_3_ (to list a few) do not contain zero.

